# Red Light-Activated Reversible Inhibition of Protein Functions by Assembled Trap

**DOI:** 10.1101/2025.03.10.642306

**Authors:** Peng Zhou, Yongkang Jia, Xuan He, Tianyu Zhang, Chao Liu, Wei Li, Zengpeng Li, Ling Sun, Shouhong Guang, Zhongcheng Zhou, Zhiheng Yuan, Xiaohua Lu, Yang Yu

**Affiliations:** Division of Life Sciences and Medicine, University of Science and Technology of China, Hefei 230026, China; Guangzhou Women and Children’s Medical Center, Guangzhou Medical University, Guangzhou 510623, China; Institute of Biophysics, Chinese Academy of Sciences, Beijing 100101, China; School of Life and Health Sciences, Hubei University of Technology, Wuhan 430068, China; Key Laboratory of Marine Genetic Resources, State Key Laboratory Breeding Base of Marine Genetic Resources, Fujian Key Laboratory of Marine Genetic Resources, Fujian Collaborative Innovation Centre for Exploitation and Utilization of Marine Biological Resources, Third Institute of Oceanography Ministry of Natural Resources, Xiamen 361005, China; Center for Reproductive Medicine, Guangzhou Women and Children’s Medical Center, Guangzhou Medical University, Guangzhou 510623, China

**Keywords:** Optogenetics, Red light, *Dr*BphP, Nanobody, LARIAT, LDB

## Abstract

Red light offers strong tissue penetration and low phototoxicity, making it an attractive option for optogenetic applications. However, the available red light-inducible optogenetic tools are rather limited. Here, we present a novel red light-induced protein clustering system, namely R-LARIAT (Red <u>L</u>ight-<u>A</u>ctivated <u>R</u>eversible Inhibition by <u>A</u>ssembled <u>T</u>rap), that controls protein functions in a spatiotemporal manner. Our system capitalizes on the rapid and reversible binding of LDBs (nanobodies-based dimerization binders) to a bacterial phytochrome *Dr*BphP, which uses a mammalian endogenous biliverdin chromophore to absorb red light. An anti-GFP nanobody fused with LDBs allows this method to quickly trap a wide variety of GFP-tagged proteins into light-induced protein clusters. Strikingly, our system exhibits an excellent performance in clustering efficiency with high light sensitivity and stability, it can function even when shielded by multiple glass plates. By utilizing the R-LARIAT system to trap and sequester tubulin, cell cycle progression can be blocked in HeLa cells. Therefore, the R-LARIAT system takes advantage of red light with greater tissue penetration and holds the potential to precisely control protein functions in living organisms.

## INTRODUCTION

Nature has evolved diverse organisms, including plants, algae, bacteria, fungi, and corals, that serve as rich sources of photoreceptors^*1, 2*^. These photoreceptors usually absorb light from 300 nm (ultraviolet; UV) to 800 nm (near-infrared light; NIR) range, triggering photochemical reactions that may induce conformational changes of photosensitive domains. Such light-induced conformational changes can be relayed to an attached effector domain to control various functions of proteins of interest (POIs) in optogenetics^*3–5*^. Optogenetic technology has been widely used to modulate protein activity with high spatiotemporal resolution, such as CRY2-CIB1^*6*^, iLID-sspB2^*7*^, LOV2^*8, 9*^ and the Magnet system^*4*^. These photosensitive modules have been engineered into proteins of interest or anchored to the plasma membrane to manipulate cellular functions, including cell dynamics^*4*^, signal transduction^*10*^, and gene expression^*11, 12*^. Additionally, optogenetic switches can also be applied to the split-protein reassembly system, allowing light control of various bioactive proteins including nucleases^*13, 14*^, recombinases^*15, 16*^, proteases^*17*^, polymerases^*18*^, antibodies^*19*^, and neurotoxins^*20*^. However, most optogenetic strategies focus on protein activation, those strategies for precise and effective inhibition of target proteins in a spatiotemporal specific manner are very limited.

Traditional genetic perturbation methods, including gene mutation or deletion and RNA interference, have been widely used to study protein function, but such genetic strategies are typically irreversible and require a relatively long time to exert their effects, and tend to induce side effects, such as lethality early in development^*21–23*^. Optogenetic tools provide a promising opportunity for inhibiting the activities and functions of proteins with rapid reversibility and high spatiotemporal resolution. UV-B photoreceptor from *Arabidopsis*, UVR8 exits as homodimers in the dark and dissociates into monomers in response to UV-B light (280–310 nm)^*24*^. Two copies of UVR8 were fused to an endoplasmic reticulum (ER)-processed protein, which led to sequestration of the fusion protein in the ER in darkness^*25*^. UV-B light induced the release of the fusion protein from the ER by disassembly of the dimers. Another versatile optogenetic strategy called LARIAT (Light-Activated Reversible Inhibition by Assembled Trap) can also inhibit protein function by reversibly sequestering target proteins into large clusters in living mammalian cells^*21*^.

The LARIAT system based on blue light (450–500 nm) is comprised of a photoreceptor cryptochrome 2 (CRY2)-fused the anti-GFP nanobody and a cryptochrome-interacting basic-helix-loop-helix1 (CIB1)-fused multimeric protein (MP; CIB1-MP). Upon blue light stimulation, the CRY2 proteins form simultaneously homo-oligomers and heterodimers with CIB1-MP, which drives the formation of large clusters through interconnections among CIB1-MP to trap and inactivate GFP-labeled proteins captured by anti-GFP nanobody^*26, 27*^. However, the limited tissue penetration capability of blue light greatly hampers LARIAT’s *in vivo* application. Since the majority of optogenetic tools based on short-wavelength lights such as violet (380–440 nm), blue (440–485 nm), or green (510–565 nm), only have limited tissue penetration capability^*8, 28, 29*^. As an alternative, red and near-infrared light (650–900 nm) offer better tissue penetration and exhibit lower phototoxicity than other visible and UV light ^*30, 31*^, which make them a promising and ideal candidate for optogenetic tools.

Phytochromes are a class of bilin-binding photoreceptors found in plants, cyanobacteria or algae, bacteria and fungi, that response to red and far-red light^*32, 33*^. Phytochromes can typically switch between the red light-absorbing Pr state and the far-red light-responsive Pfr state^*34*^. Phytochrome B (PhyB), derived from *Arabidopsis*, uses the phytochromobilin (PΦB) chromophore to absorb red light (660 nm), switching to an activated state where it can bind to phytochrome-interacting factors (PIFs). And this binding is reversible upon exposure to infrared light (720 nm)^*5, 35*^. The PhyB/PIFs system enables reversible nuclear localization of proteins, allowing light-regulated control of protein positioning in mammalian cells and zebrafish^*36*^. The cyanobacterial phytochrome 1 (Cph1) uses phycocyanobilin (PCB) as the chromophore and exhibits reversible dimer-to-monomer transition when light switches from 660 nm to 740 nm^*37*^. Cph1 was fused to neurotrophin receptor TrkB and FGFR1 to enable red light-inducible activation of RTK-mediated signaling^*37*^. Although these red light-activatable photoswitches have been developed and applied to regulate vary biological activities, they require addition of exogenous chromophores in mammalian cells, thereby limiting their application.

The bacteriophytochrome photoreceptor 1 (BphP1) derived from *Rhodopseudomonas palustris* uses biliverdin (BV), a metabolite abundantly present in mammals, as the chromophore^*38, 39*^, and can form a heterodimer with its natural binding partner RpsR2 upon 760-nm NIR light. And the heterodimer dissociates in darkness or under 660-nm red light irradiation^*32*^. However, PpsR2 can bind to the apoprotein of BphP1 irrespective of red light illumination, indicating high dark activity in BphP1/PpsR2 system^*40, 41*^, which greatly hinders its future utilization. Recently, several nanobodies were successfully selected to specifically bind to another BphP derived from *Deinococcus radiodurans* phytochrome (*Dr*BphP) with low dark activity and high specificity under 654-nm red light illumination^*40–42*^. The *Dr*BphP-nanobody heterodimers dissociate from Pfr state to fr state under 780 nm irradiation or in the darkness. Photoswitches based on *Dr*BphP and its nanobody-based binders have been used to reversibly regulate gene expression and signal pathway in living cells and mammalian animals^*40, 41*^. However, the red light-controlled reversible inactivation system by trapping or sequestering POIs is currently unavailable.

Here, we developed a versatile optogenetic strategy controlled by red light (660 nm) for reversibly inhibiting target proteins, by engineering the photosensor *Dr*BphP fused with multimeric proteins and the nanobody-based binders LDBs (Light Dependent Binders) fused with nanobodies of tagged proteins, namely R-LARIAT (Red Light-Activated Reversible Inhibition by Assembled Trap). The R-LARIAT allows rapid and reversible control of protein function by sequestering tagged proteins captured by nanobodies of tagged proteins into large clusters with red light illumination. We demonstrated that 2× LDBs connected by two copies of GGGGS-linker dramatically increase clustering efficiency for effective GFP-tagged proteins inhibition by sequestration. We also showed that the R-LARIAT has been tested for disrupting mitotic progression by trapping Tubulins. Our R-LARIAT system take advantage of red light with deeper penetration ability, and photochemical properties of *Dr*BphP that use the mammalian endogenous metabolite as chromophores, offering new application prospects for deep tissue even living animals in the future.

## RESULTS AND DISCUSSION

### Red light irradiation did not cause significant DNA damage

LARIAT is an effective strategy to reversibly inactivate proteins of interest by sequestering them into clusters with blue light^*21, 26*^. However, long-term exposure to blue light might cause DNA damage to mammalian cells, which limits the application of this technology *in vivo*^*43, 44*^. To evaluate whether red light is more advantageous in phototoxicity, we compared the effects of red light (450 nm) versus blue light (660 nm) using phosphorylated H2A.X (γ-H2A.X) staining as an indication of DNA damage in human U2OS cells (Figure 1A). After exposing the cells to either red or blue light for 2 hours, we observed a significantly increased percentage of γ-H2A.X positive nuclei under blue light exposure, but not red light (Figure 1B). The results indicate that red light may be a safer choice for optogenetically manipulating the activity of target proteins in living cells. Thus, considering that red light is capable of penetrating deeper tissues than shorter-wavelength blue light^*38, 45*^, we hypothesized that a red light-dependent LARIAT system could be more suitable for *in vivo* applications (illustrated in Figure 1C).

**Figure 1.**
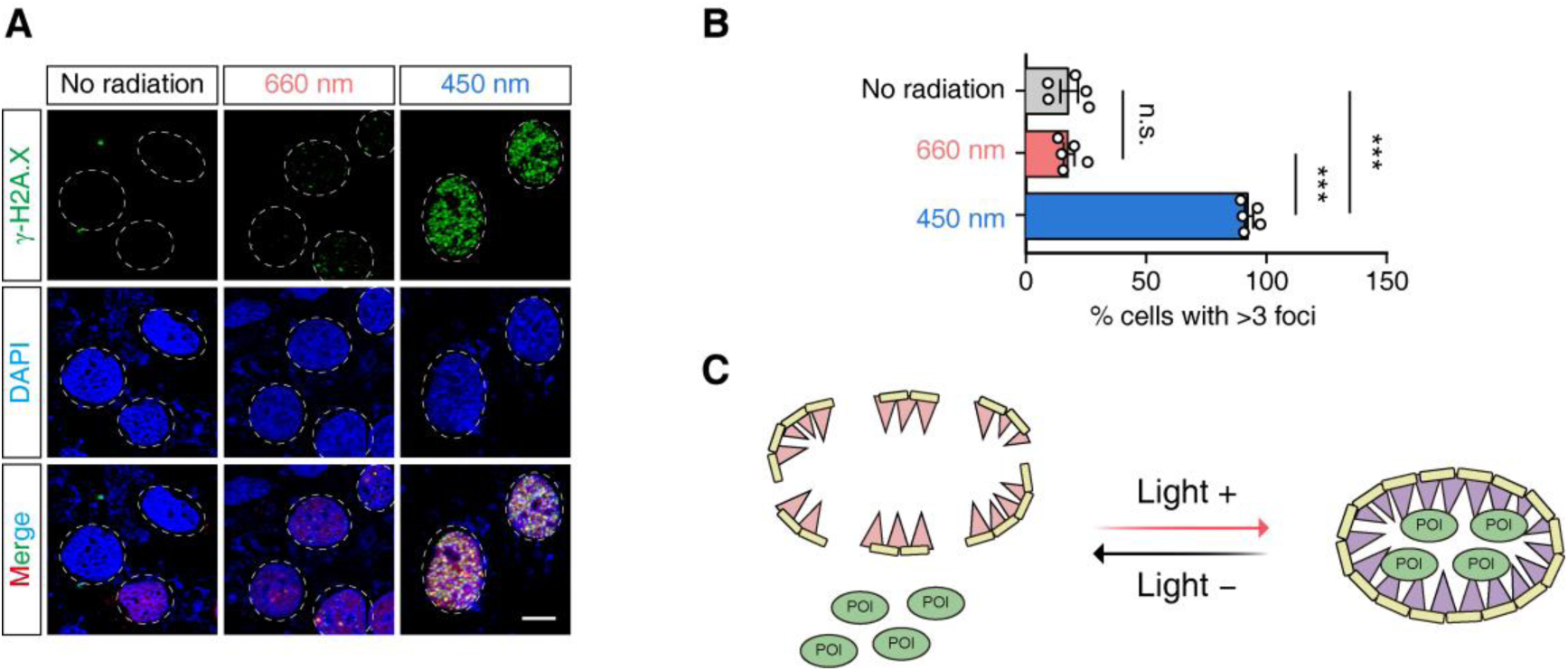
Prolonged exposure to blue light can cause DNA damage. (A) Fluorescence imaging of red light (660 nm) or blue light (450 nm)-induced γ-H2A.X expression in U2OS cells. Nuclei were labeled by DAPI. Scale bar: 10 µm. (B) Statistical data showing the percentage of γ-H2A.X foci positive cells among total DAPI-stained cells (n = 5 per group). Data are presented as mean ± SEM. Statistical analysis was performed using one-way ANOVA with Tukey’s multiple comparisons test. n.s., not significant; \*\*\**P* < 0.001. (C) Cartoon diagram showing the red light-induced LARIAT system to arrest proteins of interest (POIs).

### Creating a red light-induced protein clustering system

In the original LARIAT system^*21, 26*^, the photosensitive protein CRY2 senses blue light, triggering its oligomerization and binding to CIB1, which subsequently forms clusters to trap GFP-tagged proteins (Figure S1). Therefore, we sought to replace the CRY2 with a red light-responsive protein. A truncated bacterial photoreceptor *Dr*BphP derived from *Deinococcus radiodurans*, whose photoswitching efficiency closely matches that of the full-length *Dr*BphP^*46, 47*^, fits our needs perfectly. First, *Dr*BphP responds to a 660 nm-red light excitation wavelength that causes almost no DNA damage to cells even after prolonged exposure (Figure 1A). Second, *Dr*BphP relies on biliverdin (BV) as a chromophore, which is a natural metabolite in most eukaryotes. Third, nanobodies (i.e. LDB-3 and LDB-14) specifically recognizing and binding the red light-activated form of *Dr*BphP are readily available^*41*^.

Therefore, we cloned the *Dr*BphP and *Dr*BphP-binding nanobodies, LDBs (light-induced dimerization binders) into a single plasmid by 2A linker system to ensure constant relative amounts of *Dr*BphP and LDBs in transfected ovarian somatic cells (OSCs) of *Drosophila*. We investigated three scenarios corresponding to different designs for fusion proteins (Figure 2A and Figure S2). In scenario (I), *Dr*BphP was fused with the V_H_H(GFP) (anti-GFP nanobody) that binds specifically to GFP fusion proteins while one copy of LDB was fused with a CaMKIIα multimerization domain (MP)-mCherrry (MP-mCherry) (Figure 2A-I and Figure S2A), analogous to the blue light-dependent LARIAT (Figure S1). In scenario (II), like (I) except two copies of LDB (2× LDBs, LDB-3 and LDB-14) connected by a GGGGS linker were used (Figure 2A-II and Figure S2B). In scenario (III), 2× LDBs were fused with the anti-GFP nanobody while *Dr*BphP was fused with the MP-mCherry (Figure 2A-III and Figure S2C).

**Figure 2.**
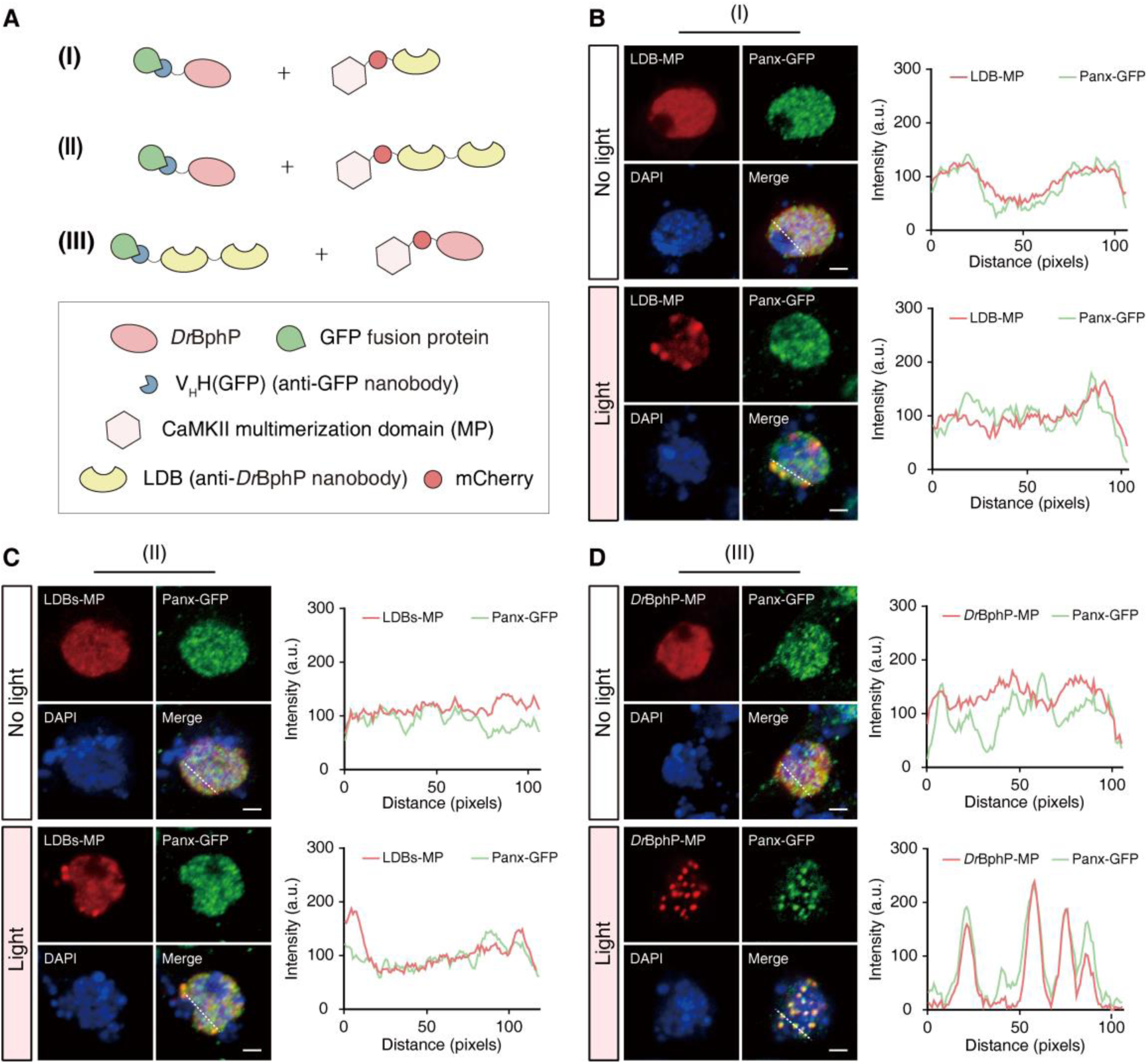
Design and validation of a novel protein clustering system induced by red light. (A) Schematic diagrams of three different scenarios for construction of fusion proteins. (B–D) Representative fluorescence images (left) and intensity profile (right) for Panx-GFP and LDB-MP (B), LDBs-MP (2× LDBs) (C), or *Dr*BphP-MP (D) in Panx-GFP cells expressing the indicated constructs as Figure 2A-I, II or III in *Drosophila* after 30 min of 660-nm illumination. Scale bar: 2 µm.

To test the ability of these constructs to form clusters with GFP fusion proteins under red light activation, we engineered a GFP knock-in tag to the C-terminus of the Panoramix (Panx) open reading frame in OSCs using CRISPR/Cas9 (Figure S3A–C). As a positive control, we tested the original LARIAT system based on CRY2/CIBN (an N-terminal fragment of the CIB1) using the Panx-GFP as a target. Consistent with the published results^*21*^, the protein clustering capability depends on wildtype CRY2 as well as the presence of both CIBN and blue light induction (Figure S4A–D). Then we tested the three constructs based on *Dr*BphP/LDBs using the same Panx-GFP cell lines with 660-nm red light illumination (Figure 2B–D). The construct (III) demonstrated the most efficient clustering ability like the blue light-dependent LARIAT (Figure S4A), with the mCherry spots (*Dr*BphP-mCherry-MP) almost completely co-clustering the GFP spots (Panx-GFP) in a red light-dependent manner (Figure 2D).

### Optimization of the red light-induced protein clustering system

Since significant clusters of Panx-GFP were observed only when LDB-3 and LDB-14 connected by a GGGGS linker peptide were fused to the C-terminus of V_H_H(GFP) (Figure 2A-III and Figure 2D). And the photosensory module (PSM) of *Dr*BphP exists as a head-to-head parallel dimer^*48, 49*^. We suspected that each nanobody module of the 2× LDBs may bind one copy of *Dr*BphP in the dimer respectively. Therefore, the distance between the two LDBs could be critical for its binding the *Dr*BphP dimer. For this, we first tested the ability of 2× LDBs with different GGGGS-linker lengths (1×, 2×, 3×, 4×) to bind *Dr*BphP by a yeast two-hybrid (Y2H) assay. Upon red light treatment, the yeast expressing *Dr*BphP and 2× LDBs with different linker lengths could grow on selective plates supplemented with 3-AT up to 10 mM (Figure S5A). Conversely, only relatively weak interactions can be detected under dark conditions (Figure S5B), indicating the interactions induced by red light between *Dr*BphP and 2× LDBs are much stronger.

Since no significant differences in strength of the interactions between *Dr*BphP and 2× LDBs with different linker lengths by the Y2H assay, we directly tested the cluster-forming ability of these different 2× LDBs in Panx-GFP cells (Figure 3A, left panel). By counting the percentage of cluster-containing cells upon red light illumination, we found that two copies of the GGGGS-linker were the most efficient constructs, in which about 70% cells with LARIAT expression could form clusters perfectly (Figure 3A, right panel and Figure 3B–3D).

**Figure 3.**
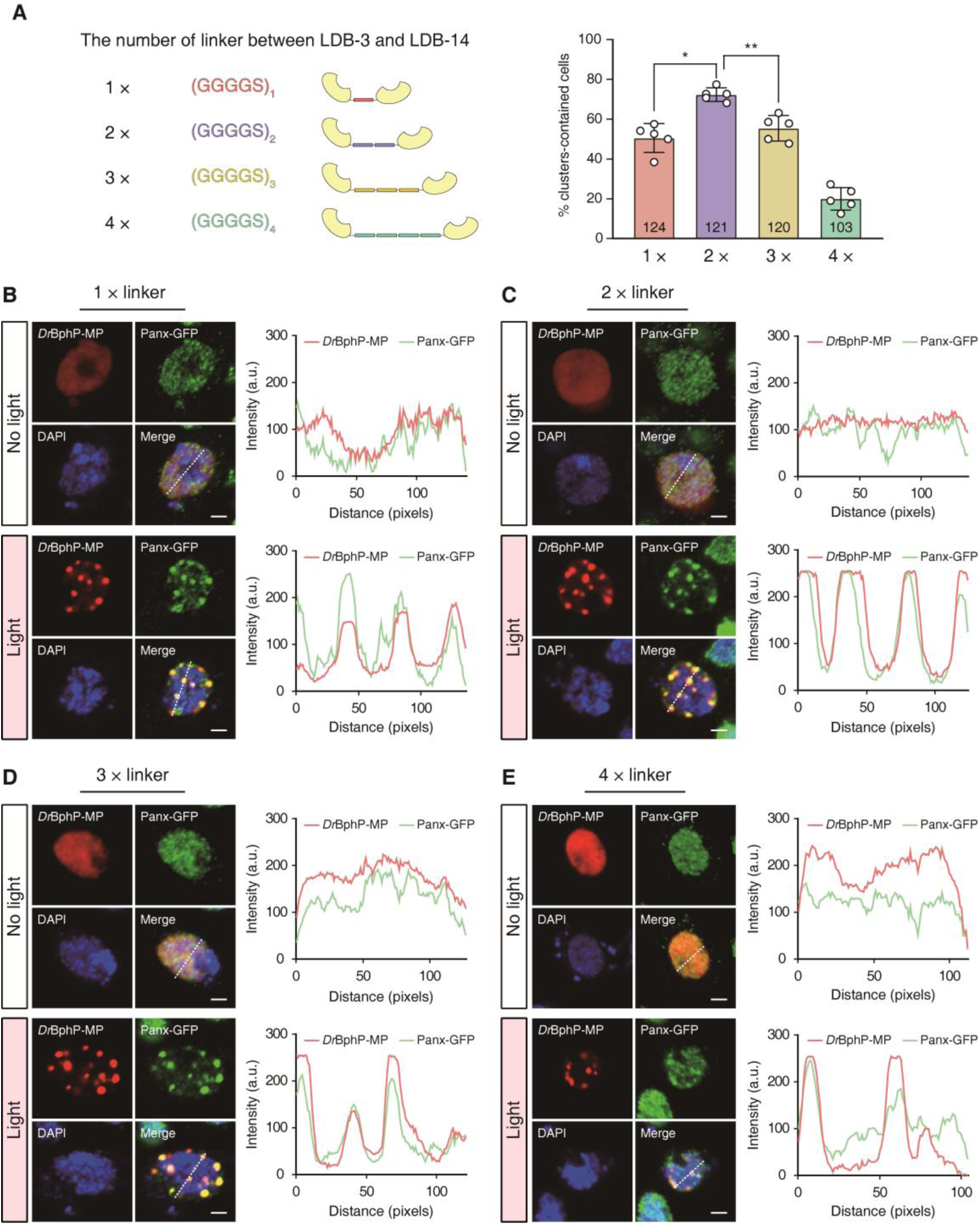
Optimization of the linker lengths in red light-induced protein clustering system. (A) Left: The sequence and length of linkers between LDB-3 and LDB-14 for red light-induced protein clustering system. Right: Statistical data of the percentage of cluster-containing cells among total mCherry positive cells after 30 minutes of 660-nm illumination (n = 5 per group). Data are presented as mean ± SEM. Statistical analysis was performed using one-way ANOVA with Tukey’s multiple comparisons test. \**P* < 0.05, \*\**P* < 0.01. (B–E) Representative fluorescence images (left) and intensity profile (right) for mCherry-MP and Panx-GFP in Panx-GFP cells transfected with the R-LARIAT plasmid containing the 1× linker (B), 2× linkers (C), 3× linkers (D), or 4×linkers (E) between LDB-3 and LDB-14 after 30 minutes of 660-nm illumination. Scale bar: 2 µm.

Therefore, the optimal clustering efficiency induced by red light was achieved when LDB-3 and LDB-14 were connected by two copies of GGGGS-linker and fused with the GFP nanobody (V_H_H(GFP)), while *Dr*BphP was fused with the MP-mCherry. Here, we present a novel red light-induced protein clustering system called R-LARIAT (red Light-Activated Reversible Inhibition by Assembled Trap) (Figure 4A-I). Similar to the original blue-light LARIAT, *Dr*BphP undergoes a conformational change upon exposure to the 660-nm red light and binds to the LDBs-(V_H_H(GFP)) fusion proteins, thereby sequestering the GFP fusion proteins into large protein clusters formed by the multimerization action of MP (Figure 4A-II).

**Figure 4.**
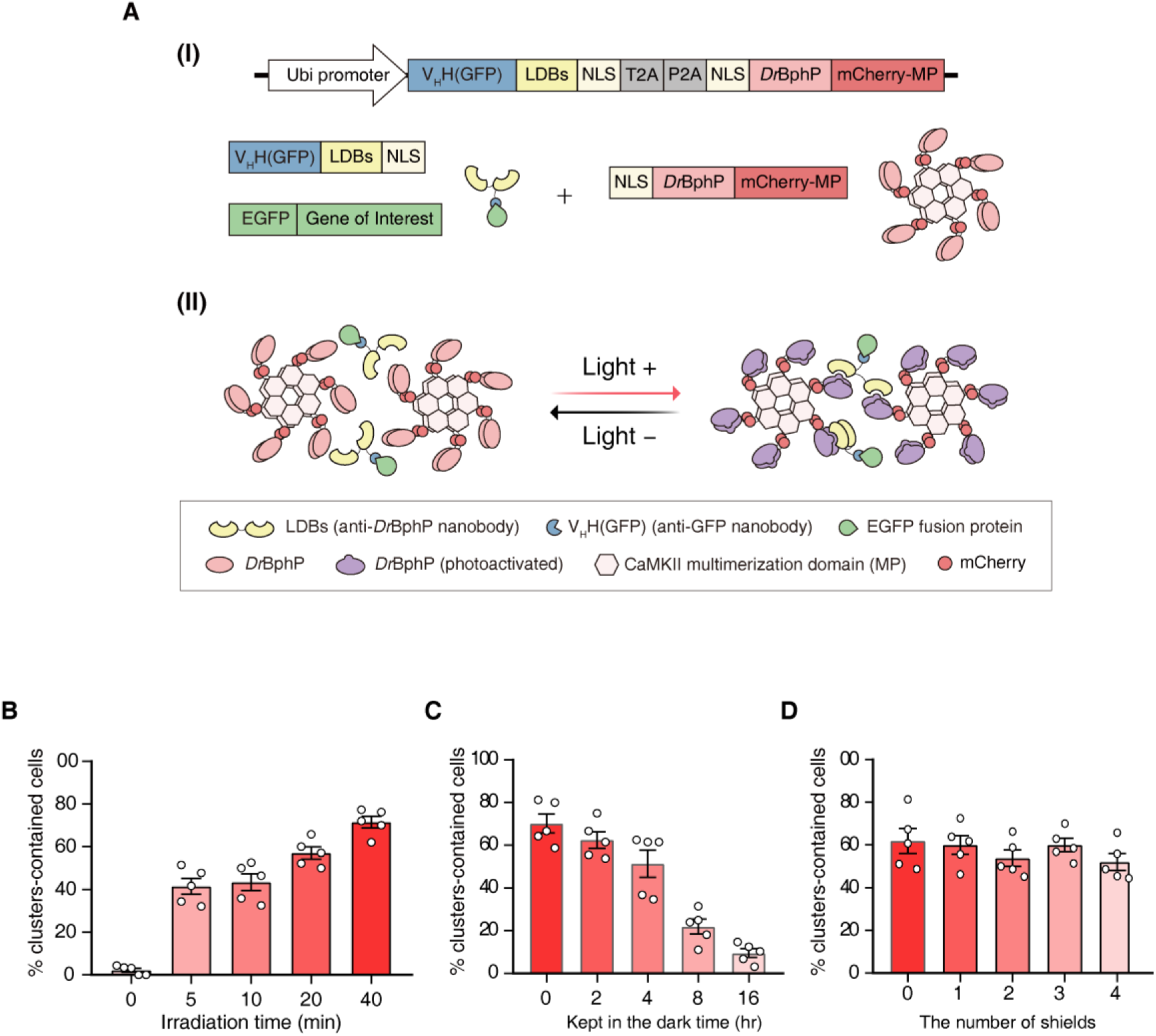
Characterization of R-LARIAT. (A) Schematic diagrams illustrating the modules (I) and working model (II) of the R-LARIAT system. The *Dr*BphP fused mCherry-MP undergoes a conformational change upon exposure to 660-nm red light and binds to the LDBs-V_H_H(GFP) fusion proteins to capture and sequester GFP-tagged proteins into large protein clusters. (B) Statistical data of the percentage of cluster-containing cells induced by red light with different irradiation time. (C) Statistical data showing the percentage of cluster-containing cells when kept in the dark at different time points after 40-min red light irradiation. (D) Statistical data showing the percentage of cluster-containing cells when illuminated through different numbers of the 1-mm thick glass plate with 660-nm red light for 20 minutes.

### Characterization of R-LARIAT

To assess the characterization of R-LARIAT, we firstly explored the time-dependence of protein clustering upon the red light exposure. We found that protein clusters could be detected in around 40% of cells expressing R-LARIAT as early as after 5-minute red light stimulation, and the percentage would reach up to approximately 70% under 40-minute of red light illumination (Figure 4B). Next, we examined the half-life of cluster dissociation by moving cells into a dark environment following a 40-minute red-light illumination. The clusters maintained at similar levels up to 4-hour darkness and then dropped significantly at 8 hours (Figure 4C). These results indicate our R-LARIAT system has considerable advantages in the aspect of cluster formation and dissociation.

Since red light is known to be able to penetrate deep tissues^*38, 45*^, we tested whether our R-LARIAT system could function well through glass plates (1 mm thickness). We placed 0, 1, 2, 3, or 4 glass plates on the top of cells and illuminated cell through the glass plates with 660 nm-red light for 20 minutes. Strikingly, even with four glass plates covering the cells, the efficiency of protein clustering remained at approximately 60% as if the glass plates were never there (Figure 4D).

To further test the ability of R-LARIAT to sequester other GFP-tagged proteins, we engineered a GFP knock-in tag at the N-terminus of the Eggless (Egg) open reading frame in OSCs (Figure S6A). We found that GFP-Egg spots almost completely co-clustered the MP-mCherry spots under red light illumination (Figure S6B), indicating our R-LARIAT system can be used to sequester diverse GFP-labeled proteins. Similarly, dramatic cluster formation could be observed in OSCs expressing another mCherry knock-in tag at Mael locus upon red light stimulation (Figure S6C and S6D). These results indicate the potential of our R-LARIAT system for sequestering a wide variety of proteins captured by different tag proteins.

### R-LARIAT successfully sequesters Tubulin to inhibit mitosis

To demonstrate our newly developed R-LARIAT can indeed manipulate protein function, we constructed stable HeLa cell lines expressing GFP-Tubulin as a target. The *Dr*BphP-MP fusion was expressed to sense the red light and the V_H_H(GFP)-LDBs fusion was used to capture the light-activated *Dr*BphP aggregates in the GFP-Tubulin cells. As a negative control, MP alone with no *Dr*BphP module was used. Upon red light illumination, the *Dr*BphP-MP signals (stained with V5 tag antibody) could colocalize GFP-Tubulin signals (Figure 5A, bottom panel), indicating that the V_H_H(GFP)-LDBs fusion successfully sequestered GFP-Tubulin proteins into the *Dr*BphP-MP clusters. In contrast, MP alone remained diffusely distributed with little or no colocalization with GFP-Tubulin (Figure 5A, top panel).

**Figure 5.**
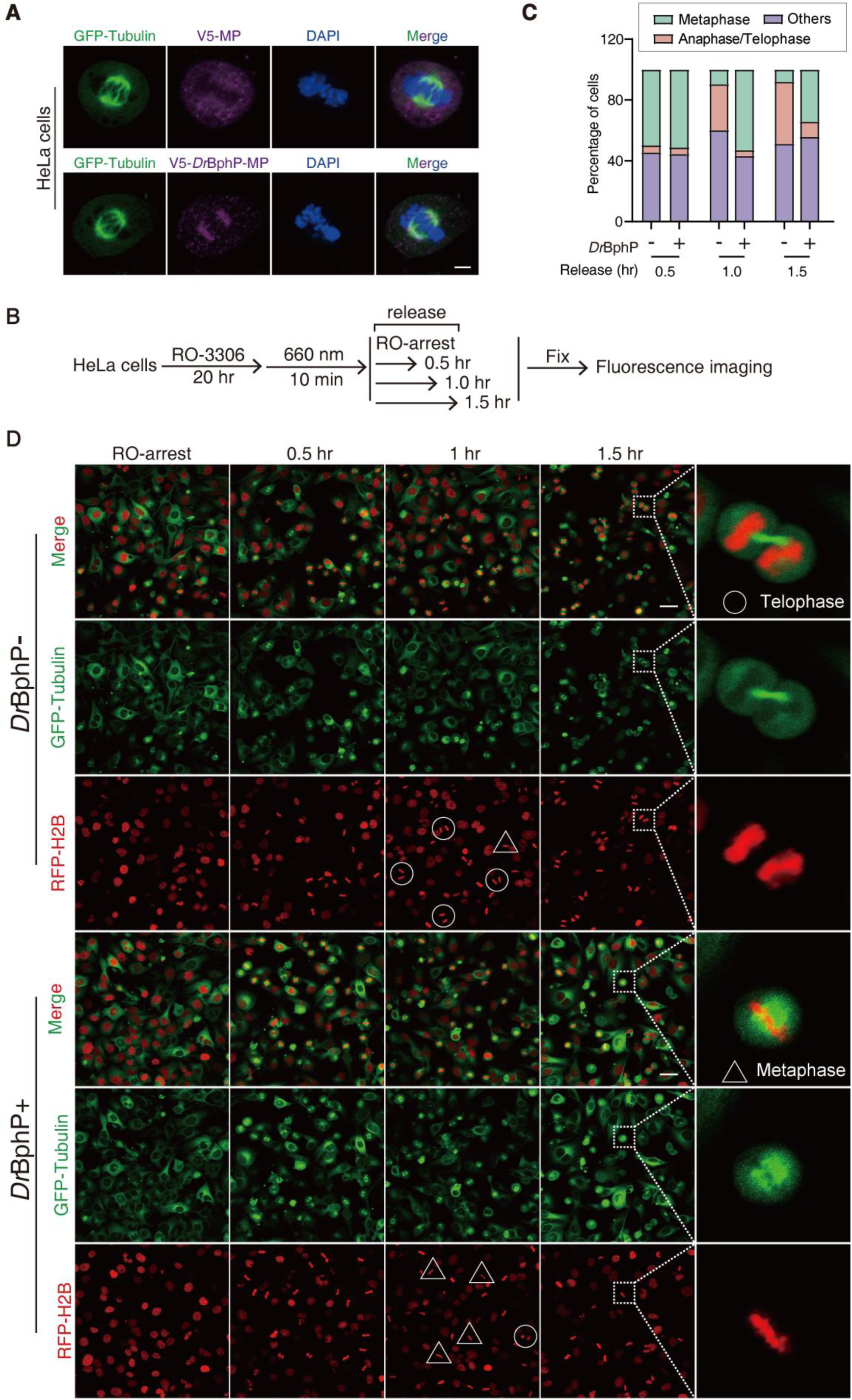
R-LARIAT can inhibit mitotic progression by trapping Tubulin. (A) Representative fluorescence images showing the colocalization of GFP-Tubulin and *Dr*BphP-MP in HeLa cells expressing R-LARIAT upon red light illumination. Scale bar: 1 µm. (B) A schematic depicting workflow of blockage of cell cycle. (C and D) The percentage of cells in different phases (C) and the expression of RFP-H2B and GFP-Tubulin (D) after withdrawal of RO-3306 for 0.5, 1 and 1.5 hours in HeLa cells expressing R-LARIAT with (*Dr*BphP +) or without *Dr*BphP (*Dr*BphP −) upon red light illumination. M phase cells were classified according to the morphologies of GFP-Tubulin and chromosomes (RFP-H2B labelled). White circles denote cells in telophase while triangles denote cells in metaphase. Scale bar: 15 µm.

To examine the functional consequences of GFP-Tubulin sequestration, we synchronized the HeLa cells with the CDK1 inhibitor (RO-3306) by arresting most of the cells at the G2/M transition. Then the cells were illuminated with 660 nm-red light for 10 min to sequester GFP-Tubulin. Then RO-3306 was washed away to allow the cell cycle to proceed. We collected cells for statistical analysis of cell cycle distribution at different time points (0.5, 1, and 1.5 hours) following the cell cycle progression (illustrated in Figure 5B). M phase cells could be classified into three categories, metaphase (highlighted with triangles); anaphase/telophase (highlighted with circles) and others, based on GFP-Tubulin and DNA morphologies (labeled with RFP-H2B)^*50*^. Consistent with the published results^*21*^, sequestering GFP-Tubulin by R-LARIAT led to a significantly increase in the percentage of metaphase cells, accompanied by a decrease in the anaphase/telophase phase after withdrawal of RO-3306 for 1 and 1.5 hours, but this phenomenon was not observed when using R-LARIAT without *Dr*BphP (Figure 5C and 5D).These results indicated that the cell cycle was significantly slowed down by the R-LARIAT trapping the GFP-Tubulin. Therefore, our newly developed R-LARIAT can be used to efficiently manipulate protein functions with high spatiotemporal resolution through light-induced clustering in living cells.

## CONCLUSIONS

Red light, with its high tissue penetration ability and low photodamage characteristics, has become one of the preferred choices in optogenetics^*31*^. However, the development of red light optogenetic tools is rather limited, restricting their application at the tissue level. To bridge this gap, we developed a novel red light-induced protein clustering system designed to overcome the limitations of existing technologies with the hope of broadening the applications of optogenetics.

Our R-LARIAT system relies on the rapid and specific interaction between the red light-sensitive *Dr*BphP and its binder nanobody LDB^*41*^. This interaction ultimately forms large protein clusters through promoting interconnection among *Dr*BphP-conjugated multimeric proteins (MPs), enabling the trapping of GFP-tagged proteins captured by a GFP-specific nanobody (V_H_H(GFP)) to eliminate their functions. Through this mechanism, we successfully constructed a red light-induced protein clustering system capable of precisely controlling the aggregation state of target proteins, thereby disturbing their functions (Figure 4A). Consequently, our system demonstrates a high light sensitivity and stability, as clusters could be induced in about of the 40% cells with only 5-min red-light illumination and were maintained for approximately 4 hours after the withdrawal of light.

Compared to the original blue light version of LARIAT, R-LARIAT based on red light almost does not cause significant DNA damage in living cells. Perhaps most importantly, R-LARIAT can penetrate through multiple glass plates (4-mm thickness), which holds the potential to be applied in live transparent animals such as *C. elegans* or Zebrafish. And our R-LARIAT system should be useful for controlling many cellular processes by trapping diverse target proteins into clusters to sequester their functions, such as gene expression, signaling pathway, cell metabolism, immune responses, neuronal activity, etc. Alternatively, the R-LARIAT can be coupled with the E3/ubiquitin/proteasome pathway to simultaneously cluster and degrade target proteins, potentially further increasing the robustness of this optogenetic tool. Our newly developed R-LARIAT provides researchers with powerful and flexible tools for modulating protein activity with high spatiotemporal resolution and reversibility, further expanding the versatility of this technique.

## MATERIALS AND METHODS

### Plasmid construction

The core modules of R-LARIAT, a truncated version *Dr*BphP without the histidine kinase domain, and the nobody-based binders for *Dr*BphP, LDB-3/LDB-14 were amplified from the NanoReD system^*41*^, then were cloned into the pENTR4 vector (Thermo Fisher Scientific, A10465). Subsequently, they were recombined into the pUbiquitin gateway vector for expression in flies and the pCAGG gateway vector for expression in mammals through LR Clonase II (Invitrogen)-mediated recombination. The sequence of CIBN (an N-terminal fragment of the CIB1) and CRY2PHR (the photolyase homology region of CRY2) or CRY2PHR (D387A) (a mutant of CRY2) in blue light-mediated LARIAT system were amplified from those vectors in published paper^*27*^. The anti-GFP nanobody (V_H_H(GFP)) and anti-mCherry nanobody sequences were cloned from a PHR-V_H_H(GFP) vector^*27*^ and a pGEX6P1-mCherry-Nanobody vector (Addgene, 70696) respectively. All primers used in this study were listed in Supplementary Table 1.

The gRNAs targeting *Panoramix* (*Panx*), *Maelstrom* (*Mael*), and *Eggless* (*Egg*) of *Drosophila* were designed using the website: http://crispor.tefor.net/, then were cloned into the CFD4 vector (Addgene, 49411), following previously described methods^*51*^. The sequence of gRNAs utilized in this research were listed in Supplementary Table 2. Gene fragments of Tubulin and H2B were amplified from the cDNA of HEK293T cells using PCR, then were cloned into the pENTR4 vector. Subsequently, pENTR4-Tubulin was recombined into the MSCV-GFP-Gateway vector through LR reaction, and pENTR4-H2B was recombined into pCW-λn-2×flag-RFP-Gateway. The plasmids information and genes sequence utilized in this research were listed in Supplementary Table 3 and Supplementary Table 4.

### Cell culture and generation of stable cell lines

The Ovarian somatic cells (OSCs) from *Drosophila*, HeLa cell line and U2OS cells used in this study originated from the Cold Spring Harbor Laboratory in the United States. The OSC cells were typically cultured in the Shields and Sang M3 Insect Media (Sigma) supplemented with 10% FBS (Fetal Bovine Serum), 5% fly extract, 0.6 mg/ml glutathione, and 10 mg/ml insulin in a constant temperature (25°C) and humidity incubator. The HeLa cells and U2OS cells were generally cultured in a DMEM media supplemented with 10% FBS and 1% Penicillin/Streptomycin in a constant temperature (37°C) and humidity (60%) incubator with 5% CO_2_.

Cells were seeded in 6-well plates to reach approximately 80% confluency for transfection. The Panx-GFP, GFP-egg and mCherry-Mael cell lines were established using CRISPR/Cas9^*52*^. The OSC cells were co-transfected with the CFD4 plasmids expressing sgRNA targeted *Panx*, *Egg* or *Mael* (1.5 µg/well), the homologous arm (1.5 µg/well) containing selective markers and inserted tags (GFP or mCherry), and the plasmid encoding wild-type spCas9 (1.5 µg/well) using FuGENE HD (Promega, E2312). The stable cell lines were generated by antibiotic selection according to the relative resistance genes at 48 hours post-transfection. The Panx-GFP and GFP-egg cell line were selected by the blasticidin (10 µg/mL), while mCherry-Mael cell line were selected by hygromycin (50 µg/mL) in OSCs. HeLa cells were co-transfected with the MSCV-GFP-Tubulin plasmid (2 µg/well) and the pCW-λn-2×flag-RFP-H2B plasmid (2 µg/well) using Lipofectamine 2000 (Invitrogen, 11668030). The stable GFP-Tubulin/RFP-H2B cell line were selected by the puromycin (1 µg/mL) and Neromycin (400 µg/mL) in HeLa cells. Then GFP-Tubulin/RFP-H2B cell line was transfected with the R-LARIAT plasmids by using Lipofectamine 2000 and selected by blasticidin (10 µg/mL) for establishment of GFP-Tubulin/RFP-H2B/R-LARIAT cell line. All the cell lines were validated via genomic PCR, RT-PCR, and Western blotting.

### Photoexcitation experiment

Panx-GFP cells were transfected with LARIAT plasmids by using FuGENE HD. Next, the cells expressing LARIAT constructs were dissociated and seeded on the glass coverslip in a 6-well plate (Corning) at 48 hours post-transfection and allowed them to adhere for at last 6 hours before imaging. Then cells adhered to the surface were either exposed to 660-nm red light (20 mW/cm²) or kept in the dark for 30 min. While the cells expressing the original LARIAT were subjected to 450-nm blue light (20 mW/cm²) for 10 min. Subsequently, the cells were fixed and subjected to immunofluorescence or imaging directly.

### Y2H assay

The Y2H assays were performed using Y187 strain. The fragments of *Dr*BphP were cloned into the pGBKT7 DNA-BD vector (Takara, Cat.#630443), while LDB-3 and LDB-14 fragments connected by 1× linker, 2× linkers, 3× linkers, or 4× linkers were cloned into the pGADT7 AD vector (Takara, Cat.#630442). The colonies containing double plasmids were selected on -Leu-Trp YSD plates and confirmed by colony PCR. To detect the protein-protein interactions under red light illumination, single colonies in serial dilutions were plated onto -Leu-Trp-His plates with varying concentrations of 3-AT. The empty BD vectors without *Dr*BphP were used as a negative control.

#### Western blot

The cells were digested by trypsin and collected into a centrifuge tube, then were washed twice with PBS, and lysed in RIPA buffer (50 mM Tris-HCl, pH 8.0, 150 mM NaCl, 1% NP-40, 1% sodium deoxycholate, 0.1% SDS, 0.1 mM DTT, PMSF and protease inhibitor cocktail). The lysates were clarified by centrifugation at 15,000g for 15 minutes and the supernatants were subjected to boil at 95°C for 5 min with Laemmli sample buffer. Then proteins were transferred to PVDF membranes after SDS-PAGE electrophoresis for immunoblotting. The antibodies used in this study include rabbit anti-EGFP (ZENBIO, 300943), mouse anti-Flag (Sigma, clone M2, F3165), mouse anti-Tubulin (invitrogen, MA5-31466), HRP-conjugated secondary antibodies (anti-rabbit, invitrogen 31460; anti-mouse, invitrogen 31430). Signals were acquired using an automatic chemiluminescence imaging system (Tanon 5200).

### Cell cycle synchronization and photoexcitation

The Cdk1 inhibitor RO-3306 was used to enable cycle synchronization of HeLa cell expressing GFP-Tubulin and RFP-H2B at M phase as described previously^*53, 54*^. The GFP-Tubulin/RFP-H2B/R-LARIAT cells were pre-arrested at the G2/M border by treatment with 6 µM RO-3306 (Selleck, S7747) for 20 h. To determine the effect of trapped Tubulin on the cell cycle, these cells were exposed to 660-nm red light (20 mW/cm²) for 10 min, and then were washed with pre-warmed PBS (37℃) containing Ca^2+^ and Mg^2+^ (PBS +) to prevent detachment of cells. After being incubated in pre-warmed medium for 0 hr,1 hr or 1.5 hr, cells were fixed with 4% formaldehyde at room temperature for 20 min. Imaging of cells was performed using a Zeiss LSM700 to distinguish mitotic sub-phases. The percentage of cells in each category among M phase was calculated according to their morphological characteristics.

#### Immunofluorescence and imaging

The cells were digested and seeded on a sterilized cover slip (22 mm × 22 mm) in a six-well plate at 60% confluency to allow cells to settle and adhere onto the cover slip. Then culture medium was removed, and the cells were washed twice with PBS and fixed with 4% paraformaldehyde at room temperature for 15–20 minutes prior to immunostaining. Then the cells were incubated with 0.5% PBST (PBS containing 0.1% Triton X-100) at room temperature for 20 minutes and were blocked with blocking buffer (0.1% PBST containing 1% BSA) at room temperature for 1 hour. Then the cells were incubated with the primary antibody against EGFP (ZENBIO, 300943), mCherry (abcam, ab125096), γ-H2A.X (CST, 9718S) or V5 (Abclone, AE089) diluted in primary antibody dilution buffer (Abclone, P0103) at room temperature for 1 hour. The cells were then incubated with the secondary antibody at room temperature in the dark for 1 hour. The secondary antibodies used in this study include goat anti-rabbit-Alexa Fluor 488 (Beyotime, A0423), goat anti-mouse-ABflo® 555 (ABclonal, AS057) and goat anti-rabbit-ABflo® 647 (ABclonal, AS060). Images were captured using laser-scanning confocal microscopy (ZIESS-LSM700) and processed with ImageJ.

## ASSOCIATED CONTECT

### Supporting Information

The Supporting Information including Figures S1–S6 and Tables S1–S4 is provided as separated PDF file.

## AUTHOR INFORMATION

### Corresponding Author

**Yang Yu** - Institute of Biophysics, Chinese Academy of Sciences, Beijing 100101, China; Guangzhou Women and Children’s Medical Center, Guangzhou Medical University, Guangzhou 510623, China.

**Xiaohua Lu** - Institute of Biophysics, Chinese Academy of Sciences, Beijing 100101, China.

**Peng Zhou** - Division of Life Sciences and Medicine, University of Science and Technology of China, Hefei 230026, China; Guangzhou Women and Children’s Medical Center, Guangzhou Medical University, Guangzhou 510623, China; Institute of Biophysics, Chinese Academy of Sciences, Beijing 100101, China.

### Author Contributions

Y.Y., X.L. and P.Z. designed this study. P.Z. carried out the experiments and analyzed the data. Y.J., T.Z., X.H. assisted in experiments. C.L., W.L., Z.L, L.S contributed materials. P.Z., S.G., Z.Z., Z.Y., X.L, Y.Y. discussed the results and drafted the manuscript. All authors have read and approved the final manuscript.

## ACKNOWLEDGMENTS

We would like to thank the Xiaobo Wang Laboratory for generously providing the original blue light-mediated LARIAT plasmids as gifts. We extend our sincere gratitude to Dr. Changmao Chen for helpful discussions and comments on this work. This work was supported in part by grants from the Ministry of Science and Technology of China (2019YFA0508903 and 2017YFA0504200 to Y.Y.), the National Natural Science Foundation of China (81921003 and 32170605 to Y.Y.). Y.Y. was additionally supported by the start-up fund from Guangzhou Women and Children’s Medical Center.

## Supplementary Information

### SUPPLEMENTARY FIGURES

**Figure S1.**
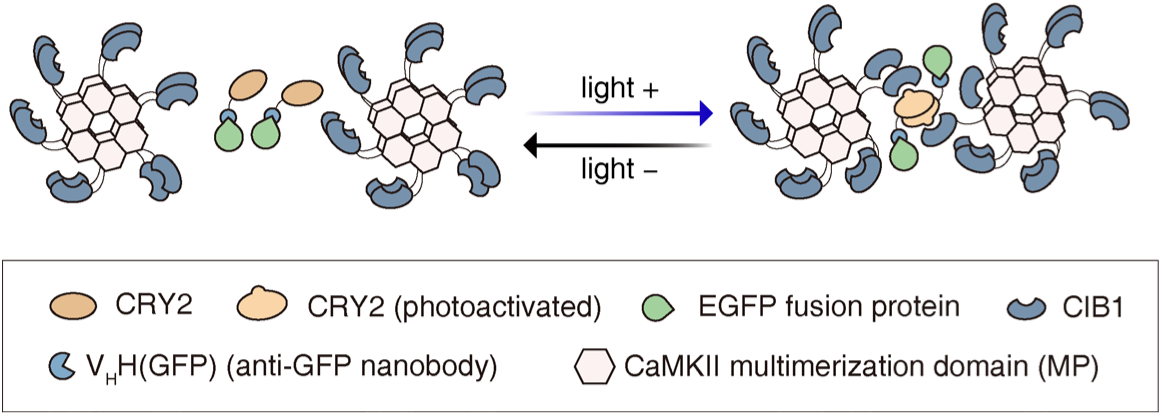
Schematic representation of blue light-induced optogenetic clustering system by LARIAT. The cryptochrome 2 (CRY2) was fused with an anti-GFP nanobody that can specifically bind to GFP-tagged proteins. The cryptochrome-interacting bHLH 1 (CIB1) was fused with the multimerization domain from CaMKIIα (MP) to form dodecamers in the cytoplasm. Blue light can trigger CRY2 oligomerization and CRY2–CIB1 dimerization, and consequently the formation of clusters to trap GFP-tagged proteins. In the dark, CRY2 reverts spontaneously to its ground state and the clusters disassemble.

**Figure S2.**
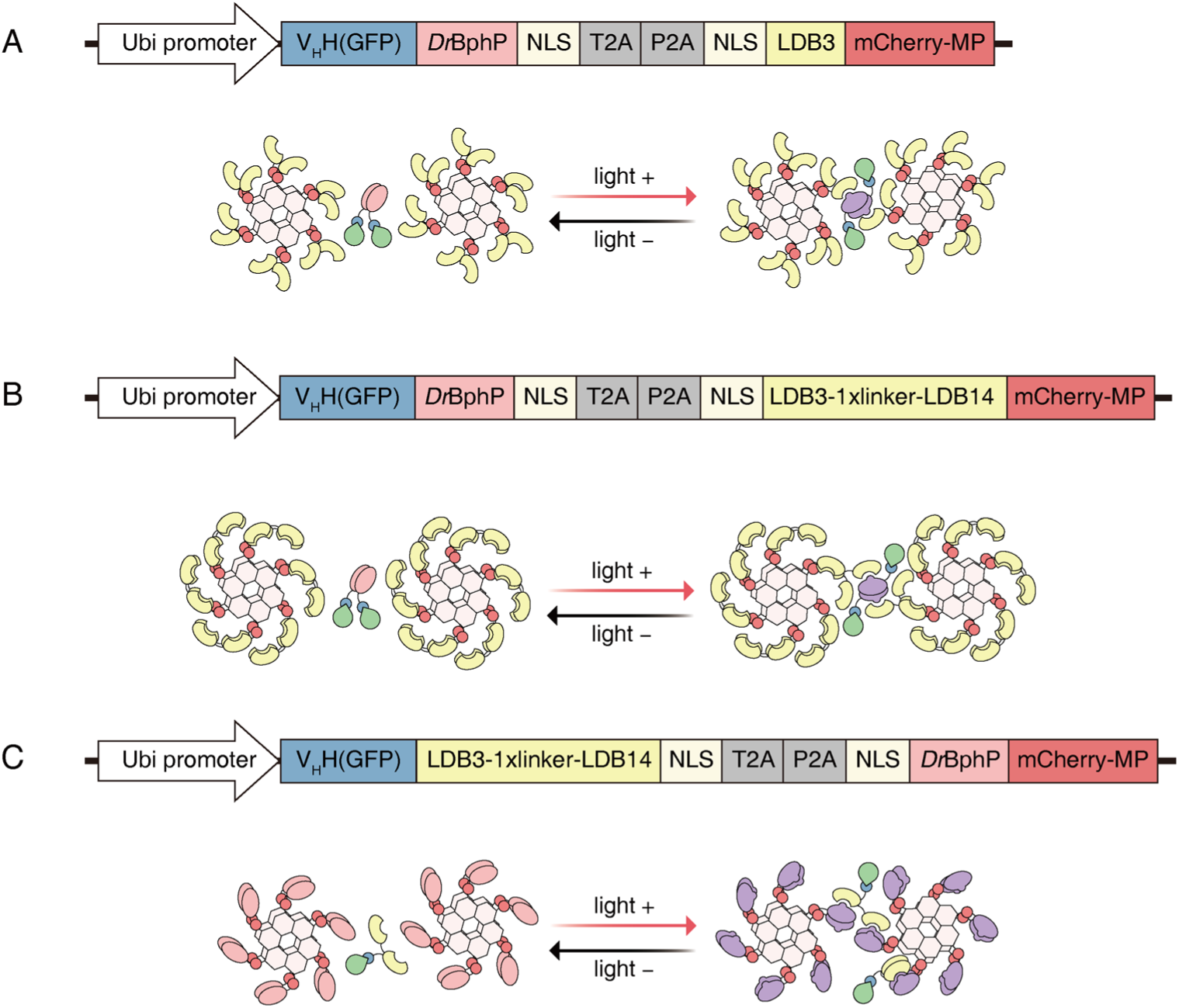
Schematic diagrams of three different constructs for protein clustering system induced by red light. (A−C) Schematic diagrams depicting construction of plasmid and protein clustering according to scenario I (A), scenario II (B) and scenario III (C). In scenario (I), The *Dr*BphP was fused with the anti-GFP nanobody (V_H_H(GFP)) while one copy of LDB (LDB3) was fused with a CaMKIIα multimerization domain (MP)-mCherrry (mCherry-MP). In scenario (II), The *Dr*BphP was fused with the V_H_H(GFP) while two copies of LDB (2× LDBs, LDB-3 and LDB-14) connected by a GGGGS linker were fused with mCherry-MP. In scenario (III), 2× LDBs connected by a GGGGS linker were fused with the V_H_H(GFP) while *Dr*BphP was fused with the mCherry-MP.

**Figure S3.**
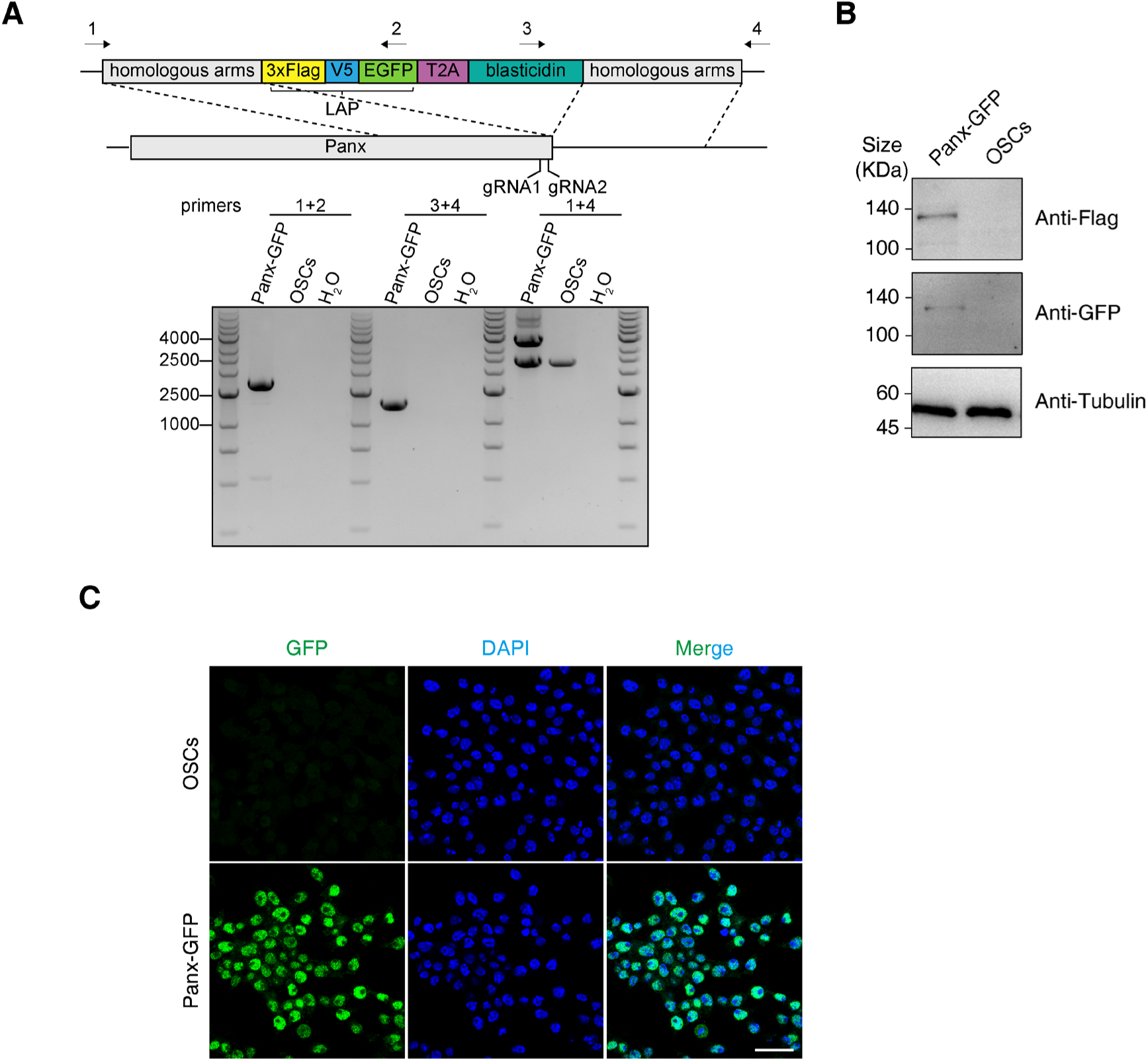
Construction of Panx-GFP cell line. (A) A schematic of showing the knock-in strategy to construct Panx-GFP cell line in OSCs using CRISPR/Cas9. Agarose gel showing PCR results confirming a correct integration of GFP-tag (3× Flag, V5 and EGFP) to the C-terminus of the Panx locus in OSCs of *Drosophila*. PCR primers amplifying the indicated regions are shown on the top while two gRNAs targeting near the stop codon of Panx are indicated at the bottom. (B) Western blots showing the expression of GFP and 3× Flag in Panx-GFP cells or OSCs. Tubulin was employed as a loading control. (C) Immunofluorescence showing the GFP fluorescence signal in Panx-GFP cells and OSCs. Scale bar: 20 µm.

**Figure S4.**
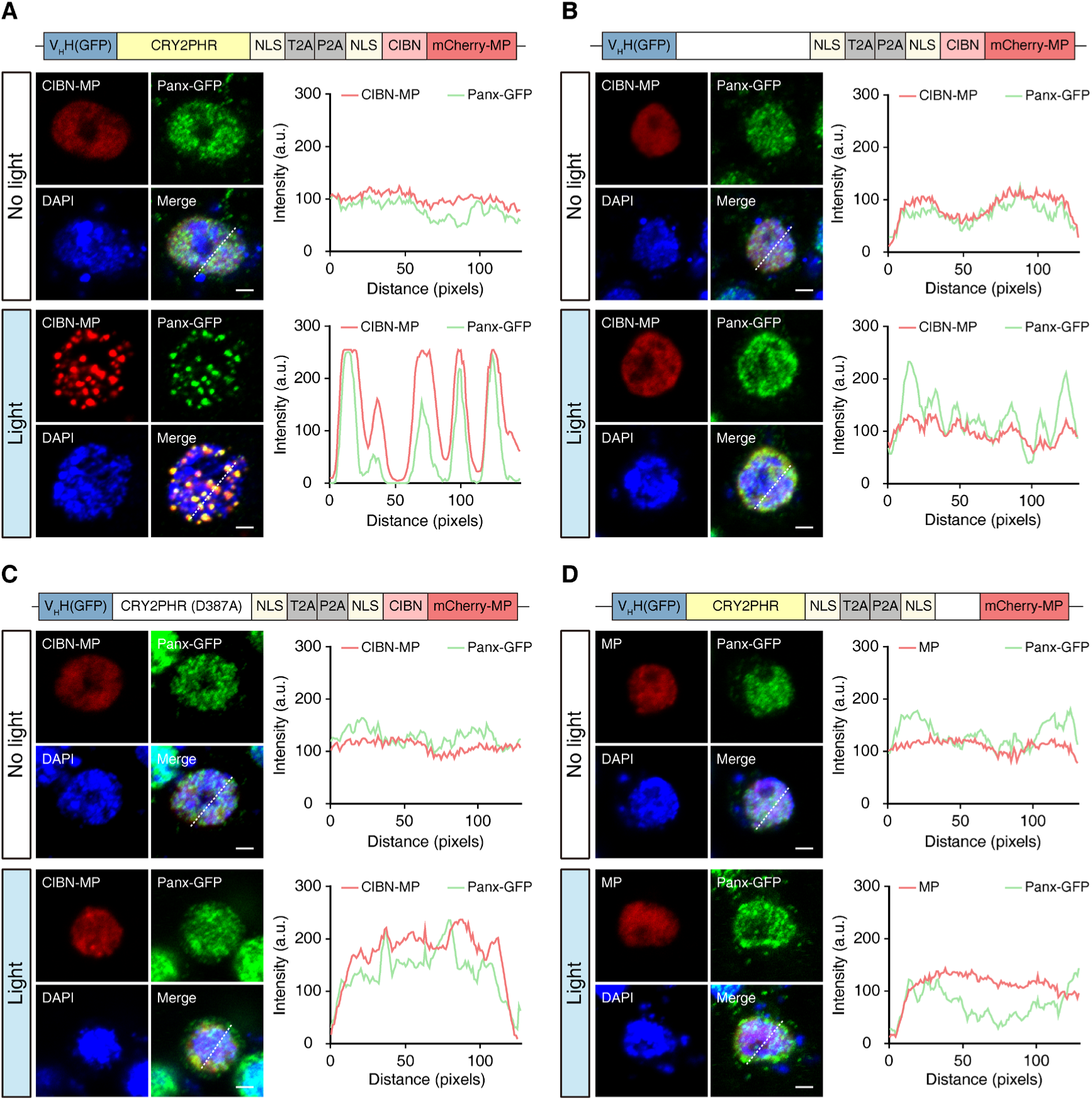
Design and validation of LARIAT modules for blue light-induced cluster formation. (A–D) Schematic diagrams of plasmid construction (top) and corresponding representative fluorescence images (left) and intensity profiles (right) for CIBN-MP and Panx-GFP in Panx-GFP cells expressing the indicated LARIAT module after 10 minutes of 450-nm blue light illumination. In Figure (A), The cryptochrome photolyase homology region (CRY2PHR) was fused with the anti-GFP nanobody (V_H_H(GFP)) while an N-terminal fragment of the CIB1 (CIBN) was fused with a CaMKIIα multimerization domain (MP)-mCherrry (mCherry-MP). In Figure (B), The V_H_H(GFP) without CRY2PHR, and CIBN fused with mCherry-MP, were cloned into a single plasmid by 2A linker system. In Figure (C), A mutant of CRY2, CRY2PHR (D387A) was fused with the V_H_H(GFP) while CIBN was fused with the mCherry-MP. In Figure (D), The CRY2PHR fused with V_H_H(GFP), and mCherry-MP without CIBN, were cloned into a single plasmid by 2A linker system. Scale bar: 2 µm.

**Figure S5.**
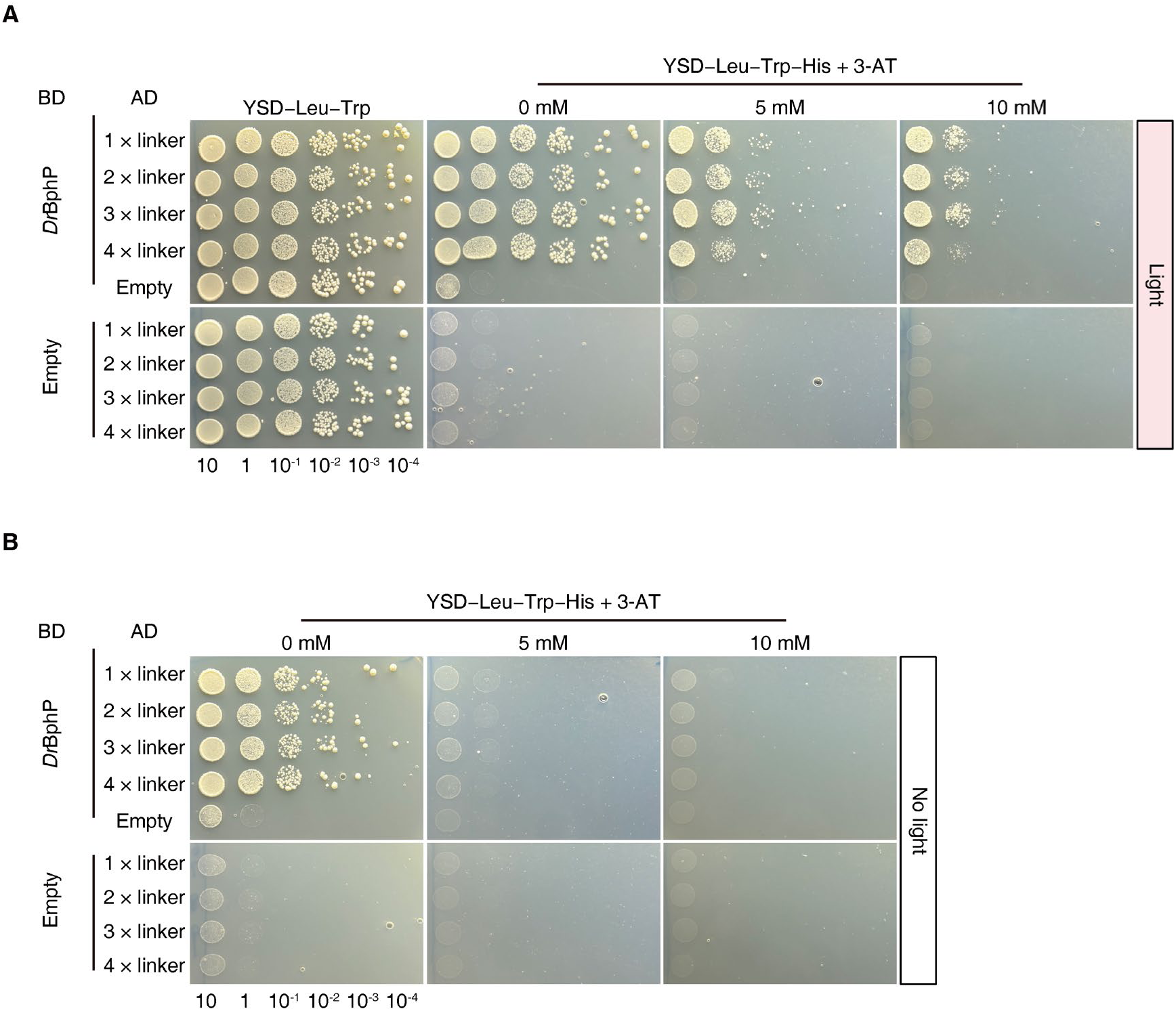
Interactions between *Dr*BphP and 2× LDBs. (A and B) Yeast two-hybrid assays showing interactions between *Dr*BphP and 2× LDBs (LDB-3 and LDB-14) with different linker lengths under red light illumination (A) or dark conditions (B). The empty BD vector without *Dr*BphP was used as a negative control. The colonies expressing *Dr*BphP and 2× LDBs were grown on YSD/–Leu/– Trp/–His medium supplemented with 0–10 mM of the 3-AT.

**Figure S6.**
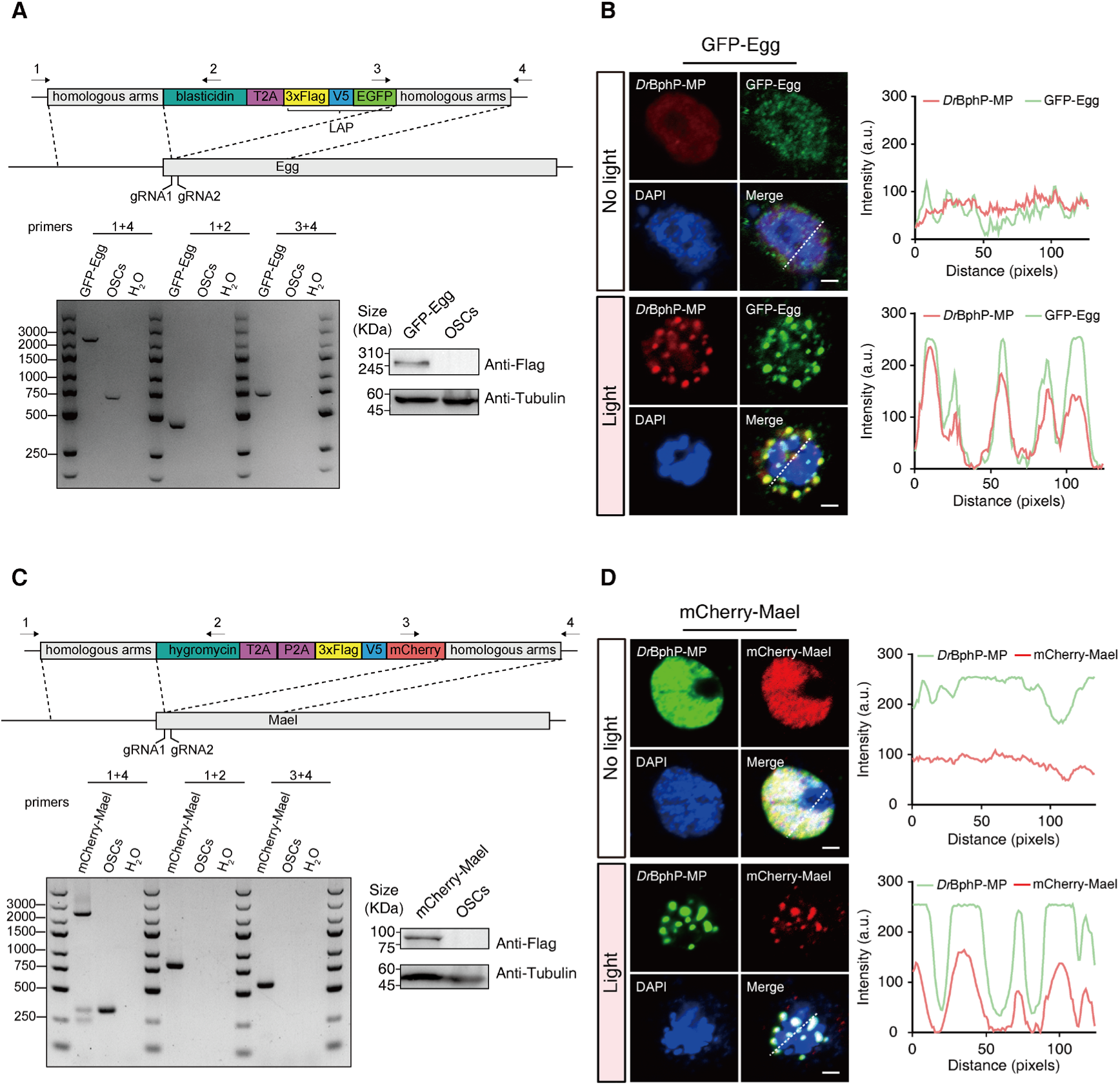
Red light-induced cluster formation in GFP-Egg and mCherry-Mael cells. (A) Schematic showing the knock-in strategy to construct Egg-GFP cell line in OSCs using CRISPR/Cas9. Agarose gel showing PCR results confirming a correct integration of GFP-tag to the N-terminus of the Egg locus in OSCs of *Drosophila*. Western blots showing the expression of 3× Flag in GFP-Egg cells or OSCs. (B) Representative fluorescence images and intensity profiles of *Dr*BphP-MP and GFP-Egg in GFP-Egg cells expressing the R-LARIAT plasmid under red light illumination or in darkness. Scale bar: 2 µm. (C) Schematic showing the knock-in strategy to construct mCherry-Mael cell line. The agarose gel and western blots showing the integration of mCherry-tag to the Mael locus in OSCs and the expression of 3× Flag in mCherry-Mael cells respectively. (D) Representative fluorescence images and intensity profiles of GFP-MP and mCherry-Mael in mCherry-Mael cells expressing the R-LARIAT plasmid. Scale bar: 2 µm.

### SUPPLEMENTARY TABLES

**Table S1.**
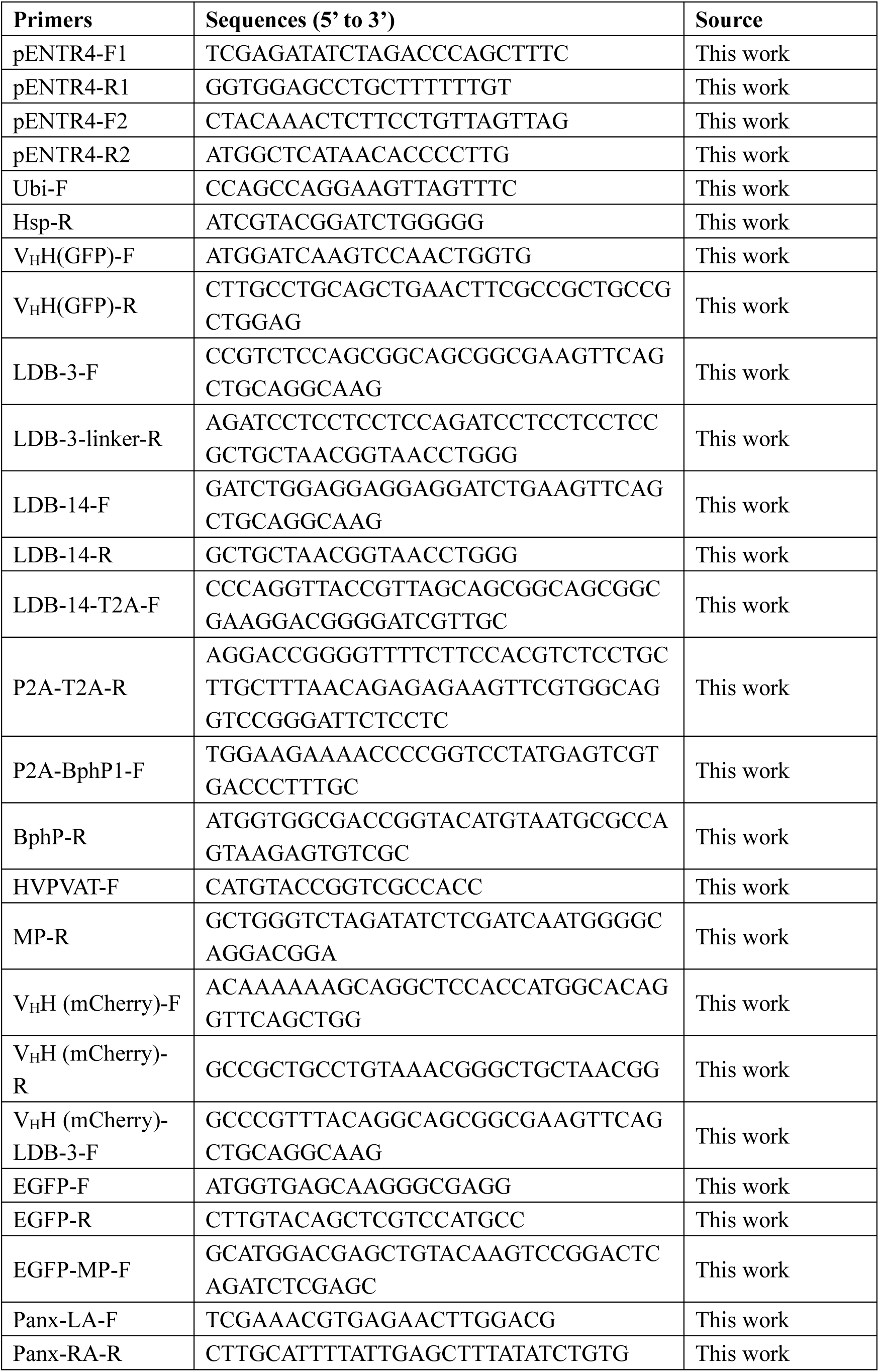
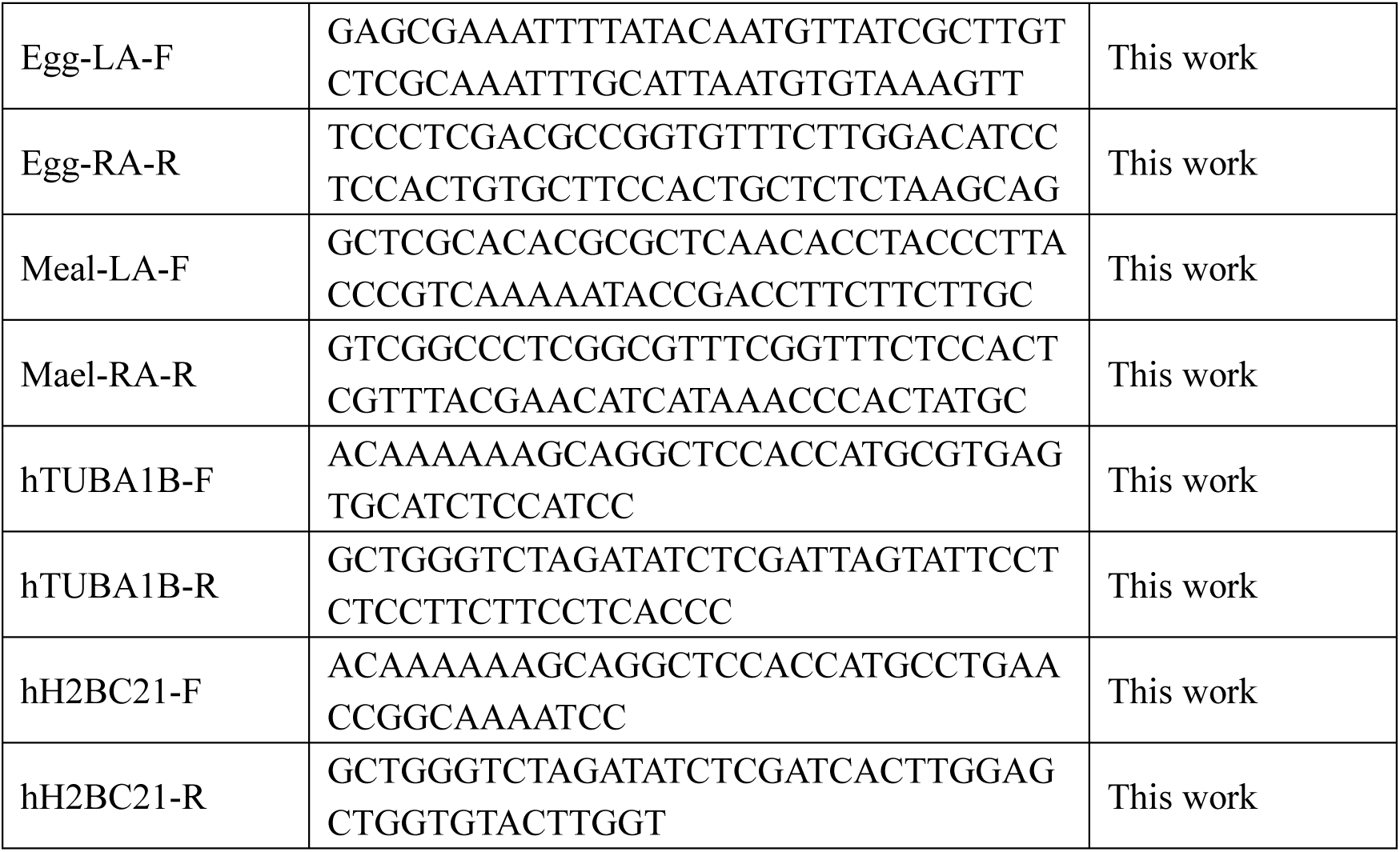
Primers used in this study.

**Table S2.**
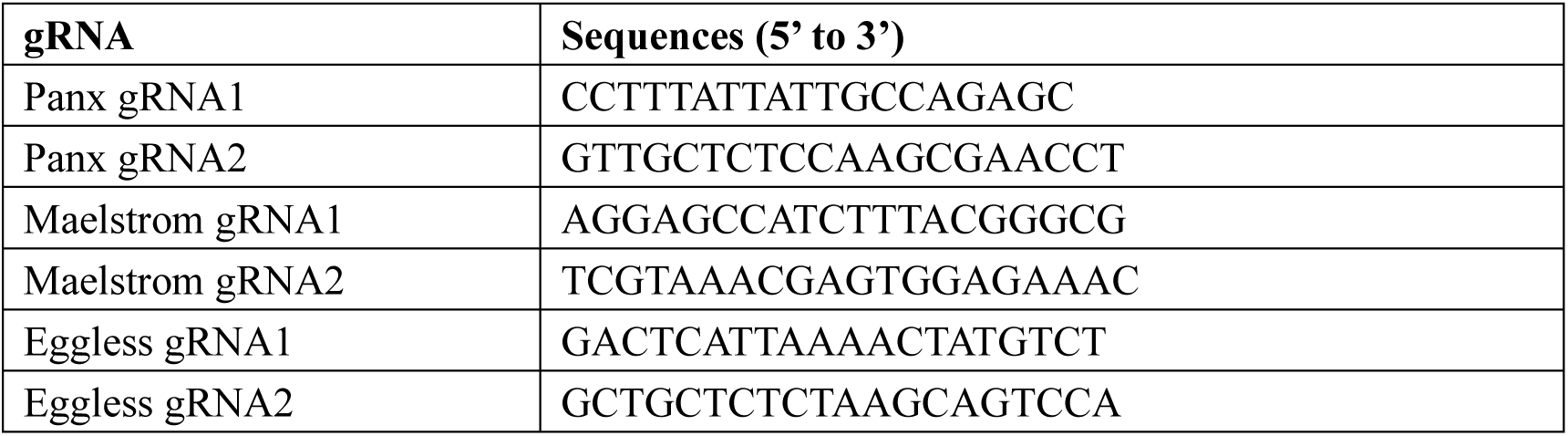
gRNA sequences for knock-in in OSCs.

**Table S3.**
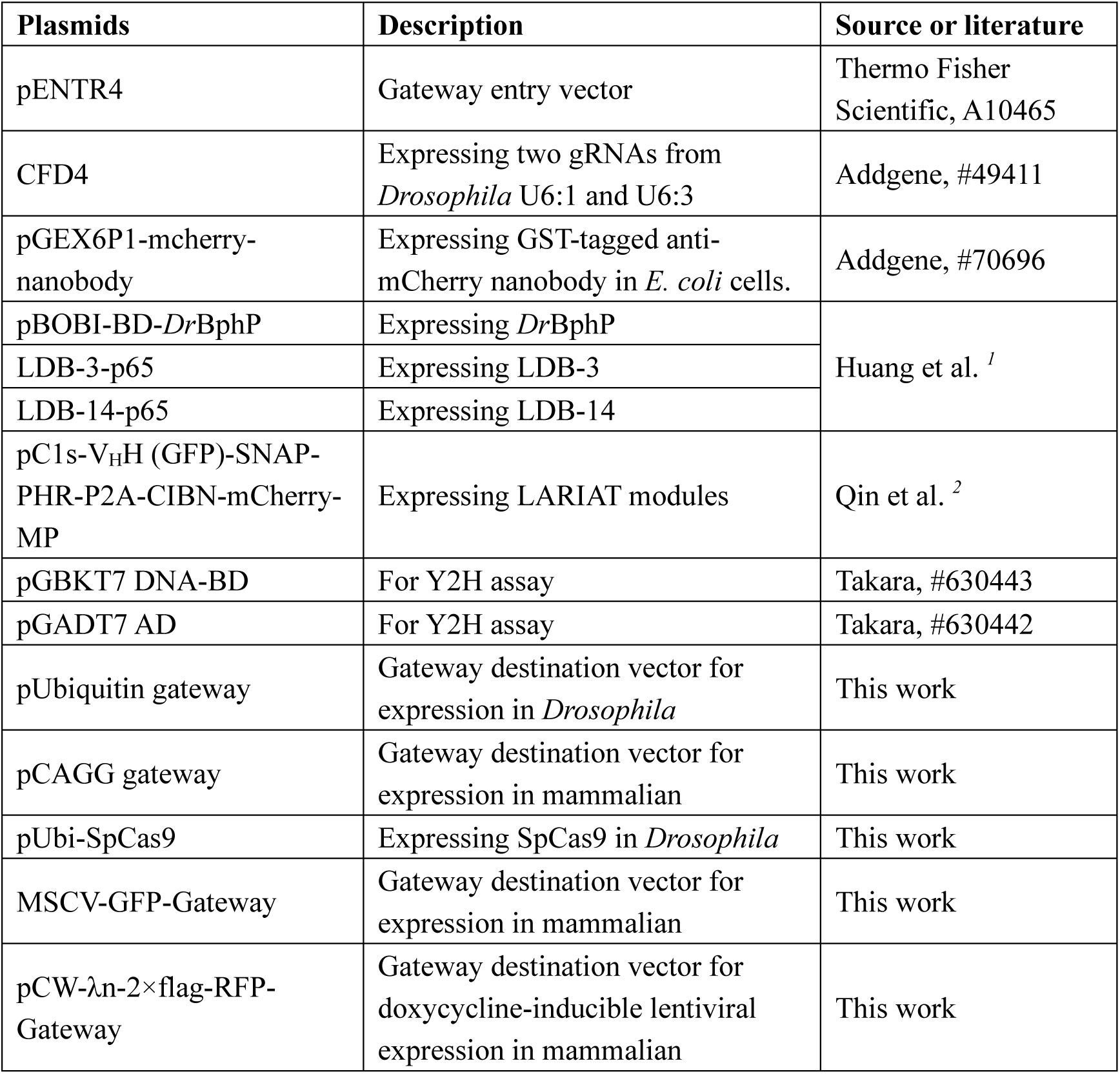
Plasmids information used in this study.

| Plasmids | Description | Source or literature |
| --- | --- | --- |
| pENTR4 | Gateway entry vector | Thermo Fisher Scientific, A10465 |
| CFD4 | Expressing two gRNAs from *Drosophila* U6:1 and U6:3 | Addgene, #49411 |
| pGEX6P1-mcherry-nanobody | Expressing GST-tagged anti-mCherry nanobody in *E. coli* cells. | Addgene, #70696 |
| pBOBI-BD-*DrBphP* | Expressing *DrBphP* | Huang et al. <sup>1</sup> |
| LDB-3-p65 | Expressing LDB-3 |  |
| LDB-14-p65 | Expressing LDB-14 |  |
| pC1s-V<sub>H</sub>H (GFP)-SNAP-PHR-P2A-CIBN-mCherry-MP | Expressing LARIAT modules | Qin et al. <sup>2</sup> |
| pGBKT7 DNA-BD | For Y2H assay | Takara, #630443 |
| pGADT7 AD | For Y2H assay | Takara, #630442 |
| pUbiquitin gateway | Gateway destination vector for expression in *Drosophila* | This work |
| pCAGG gateway | Gateway destination vector for expression in mammalian | This work |
| pUbi-SpCas9 | Expressing SpCas9 in *Drosophila* | This work |
| MSCV-GFP-Gateway | Gateway destination vector for expression in mammalian | This work |
| pCW-λn-2×flag-RFP-Gateway | Gateway destination vector for doxycycline-inducible lentiviral expression in mammalian | This work |

**Table S4.**
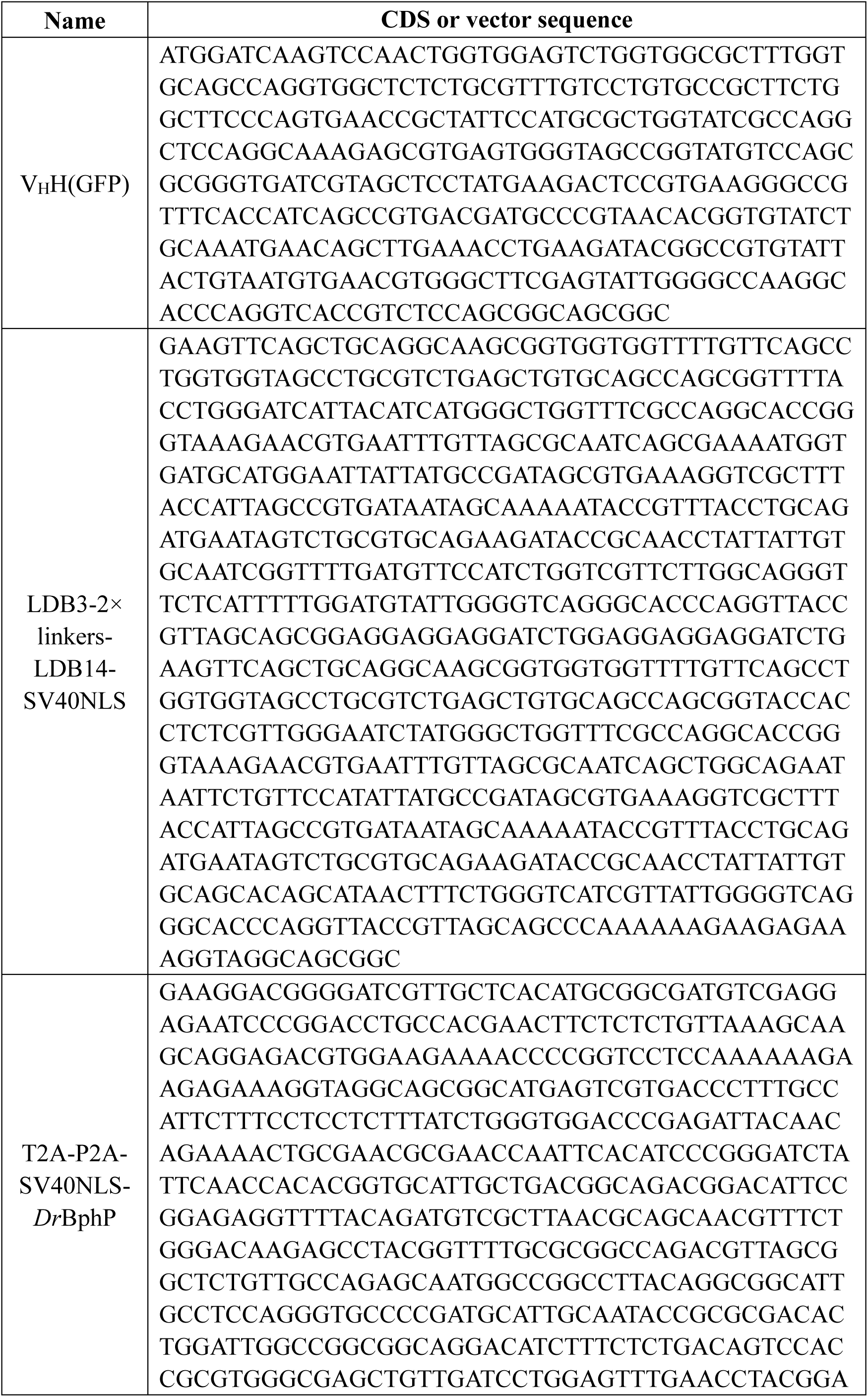
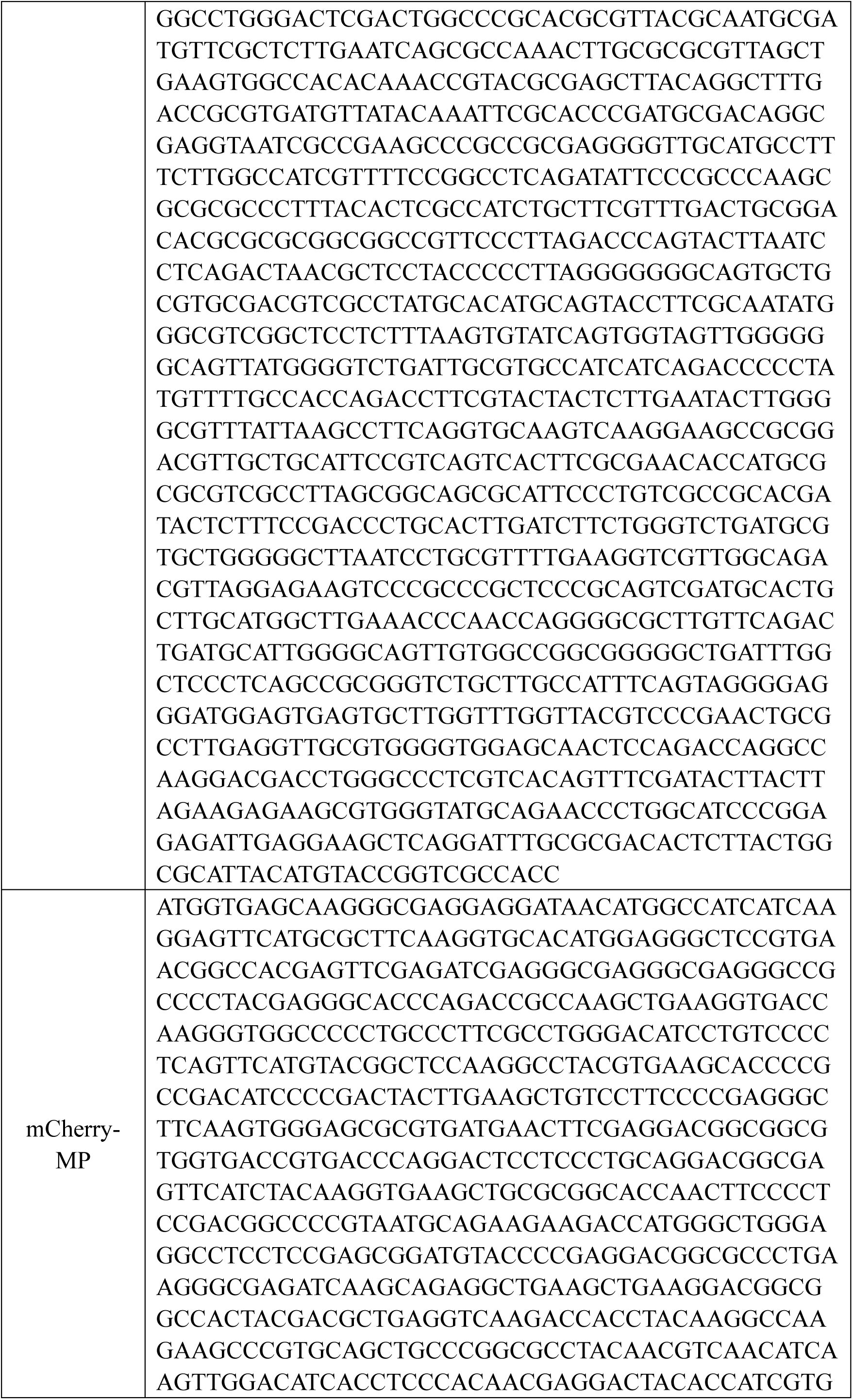
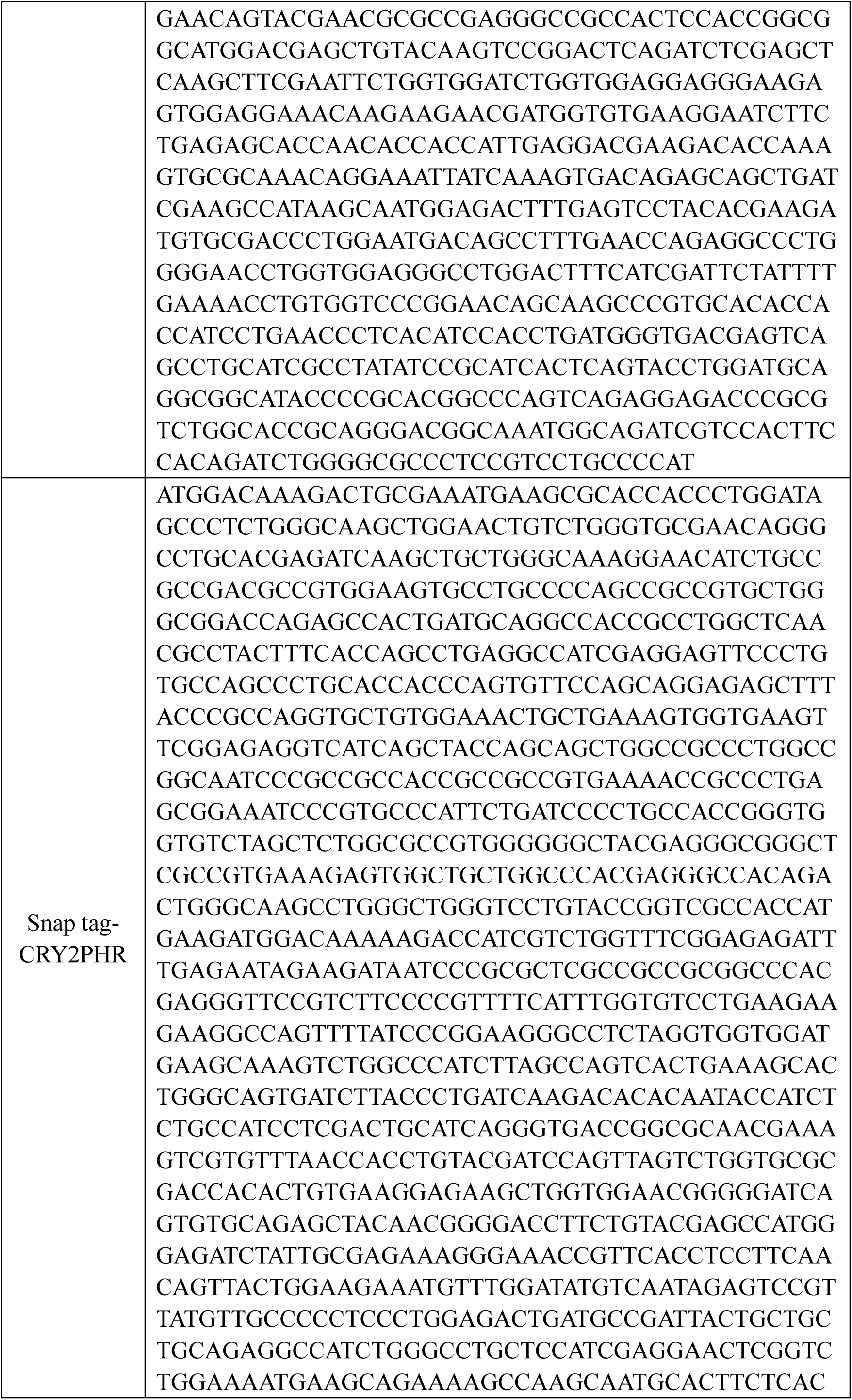
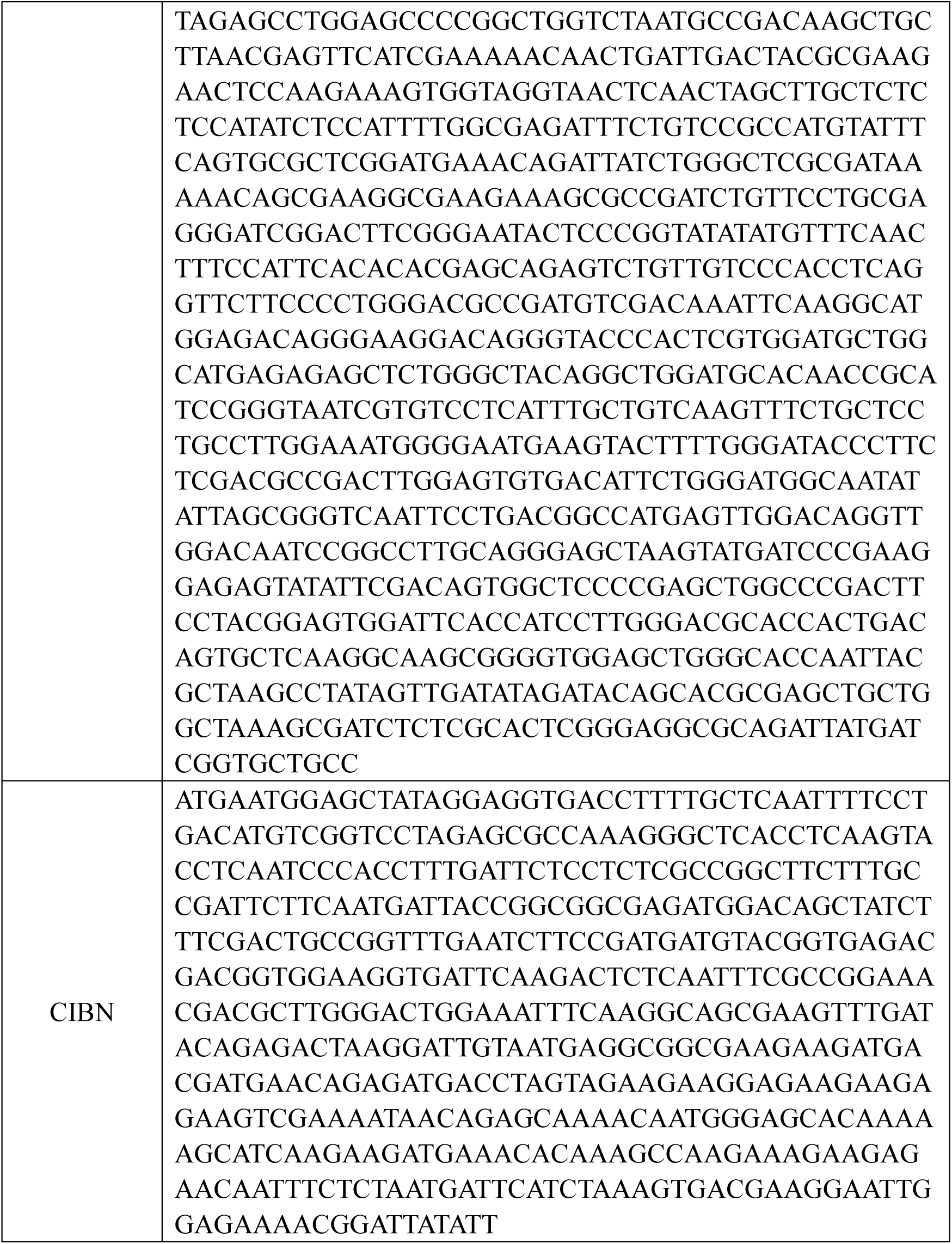
The gene sequence of LARIAT and R-LARIAT modules.

